# A personalized map of where, when, and how to stimulate the brain to elicit controlled responses

**DOI:** 10.64898/2026.07.16.738930

**Authors:** Giovanni Rabuffo, Irene Acero-Pousa, Tomas Berjaga-Buisan, Gustavo Deco

## Abstract

Brain stimulation has transformed the treatment of several neurological and psychiatric disorders and is now widely used to read out causal interactions in the human brain. However, its effects vary from one trial to the next, even when the stimulation parameters are kept identical. Identifying the sources of this variability and developing new strategies to reduce it are key to achieving more controllable interventions. Yet, exhaustive testing in real experiments is unfeasible, because each participant can only be probed at a handful of sites under conditions the experimenter cannot fully control. Here we use personalized brain models to map how stimulation responses vary across target sites and with the brain’s ongoing state. We show that stimulation responses are jointly determined by the target’s position along the cortical unimodal-to-transmodal hierarchy and the brain’s global ongoing activity, with lower-activity states yielding larger responses. Accordingly, timing stimulation to low-activity periods reduces trial-to-trial variability in response magnitude. Even without state information, joint stimulation of specific region pairs reduces variability compared with stimulating either region alone. Together, these findings identify where, when, and how to stimulate the brain to achieve more reproducible responses, with concrete predictions for closed-loop and circuit-level stimulation protocols.

## 1 Introduction

The ability to perturb the human brain directly, rather than merely observe it, has made brain stimulation a central tool in both basic neuroscience and clinical medicine. Perturbing a region and recording the response provides a direct way to map the causal organization of the brain [1–3]. In the clinic, brain stimulation is used to treat conditions such as major depressive disorder, obsessive-compulsive disorder, Parkinson’s disease, and chronic pain [4, 5]. In both research and clinical settings, outcomes remain variable. Even for an established indication such as treatment-resistant depression, only a fraction of patients reach remission, and the magnitude and time course of improvement vary widely across individuals [6–8]. Making stimulation more reliable is a shared goal of mechanistic and therapeutic work.

A basic obstacle to reliability is that the response to stimulation is not fixed. The same target, stimulated with the same parameters, evokes responses that differ several-fold from one trial to the next [9–13]. This variability is not all noise: a growing body of work shows that it depends on the brain’s ongoing activity at the time of stimulation [14, 15]. The phase of ongoing oscillations in the target region can double or halve the size of an evoked response [16–19], and broader physiological and cognitive states can also shape stimulation outcomes [5, 15, 20, 21]. This dependence is not limited to the activity of the target region. Large-scale brain dynamics measured immediately before stimulation predict the magnitude and topography of the response [3, 22–25]. Consistent with this, chronometric TMS-fMRI shows that the downstream network effect of a fixed prefrontal target also depends on the ongoing brain state [26]. The response to a pulse therefore depends on the global brain state at the moment of stimulation, not on the activity of the target region alone [15].

Despite this evidence, the spatial and temporal dependence of stimulation effects has not been characterized systematically, for two reasons. First, the experiment that would settle this question—stimulating every region, at many times, against many ongoing states, in the same participant—is infeasible: a session permits only a handful of sites, and the experimenter cannot freely set or exhaustively sample the ongoing whole-brain state at the moment of the pulse [27]. Second, even when a response is measured, the brain’s ongoing activity is strongly autocorrelated, and the post-stimulation state still carries the signature of the pre-stimulation activity [28–30]. This baseline leakage confounds the measured effect. Averaging over many trials suppresses it and reveals the mean response, but at the cost of the single-trial, state-dependent variation that is the object of interest [31].

Computational models offer a way around these limits. A model can be stimulated at every region and at every moment of its ongoing activity [32, 33], free of the limits of an empirical session. And because the same model can be run with and without the pulse from an identical starting state, the baseline leakage that confounds real measurements can simply be subtracted away. Connectome-based biophysical models offer a mechanistic understanding of how stimulation or lesions reshape brain dynamics [2, 34–36]. However, they rely on explicit model assumptions and must be re-simulated from scratch for every target and initial condition, making exhaustive sweeps computationally costly. To overcome these limitations, we adopt a data-driven framework recently introduced by Luo and colleagues [37], which captures the dynamics of an individual brain by training an artificial neural network (ANN) on empirical resting-state fMRI. Once trained, the model can be stimulated at any region and any time at negligible cost. Here we build on this framework, training personalized models of 100 participants from the Human Connectome Project (HCP), and extending it in three directions. First, we provide new external validation of the framework by showing that simulated spontaneous activity reproduces the empirically observed resting-state network dynamics beyond static functional connectivity (FC), and that the activation patterns evoked by seven cognitive HCP tasks are recovered in the model following virtual stimulation of task-relevant targets. Second, we characterize how the response to stimulation depends on where and when it is applied. Third, we use the same models to design two strategies for reducing response variability beyond what can be achieved with state-naive single-target stimulation: state-informed closed-loop stimulation and state-naive circuit-level bifocal stimulation.

## 2 Results

To map how the brain responds to stimulation across regions and brain states, we built personalized models of resting-state fMRI dynamics and stimulated them in silico. For each participant, we trained a feedforward ANN to predict the whole-brain activity vector at the next time point from the current one (Figure 1A). The training was performed on fMRI data parcellated into 400 cortical and 50 subcortical regions, so subcortical activity shapes the learned dynamics, while all perturbation analyses focus on cortical targets.

**Figure 1:**
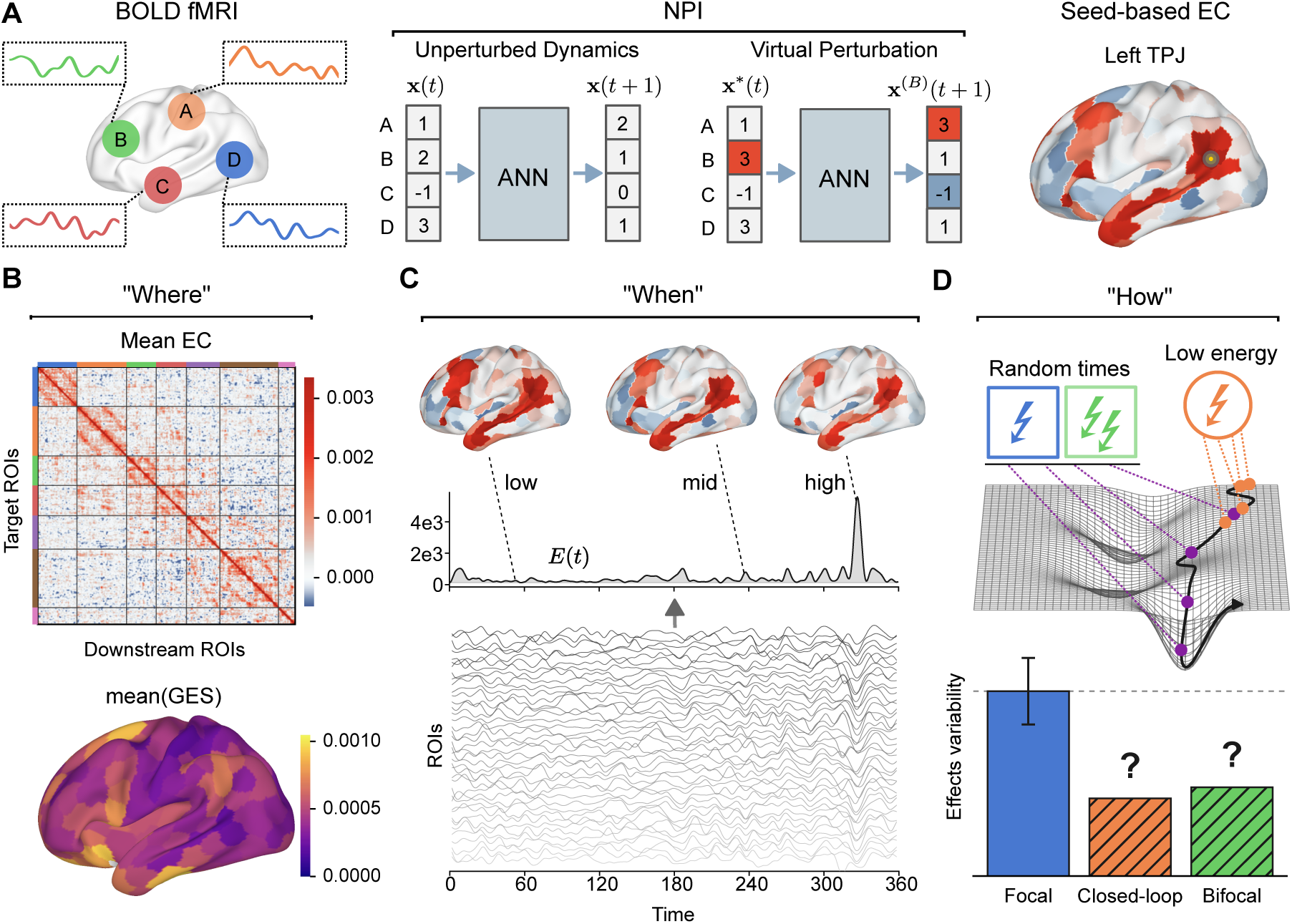
Personalized brain models of resting-state fMRI enable virtual perturbation across targets, times and brain states. **(A)** For each of 100 HCP participants, a feedforward ANN is trained to predict the whole-brain activity at the next time point from the current one, using only that participant’s resting-state fMRI; this trained network is the personalized brain model, and its output is the unperturbed dynamics. A virtual perturbation adds a fixed increment to one region (red) and propagates it through the same network; the difference between the perturbed and unperturbed predictions, over all regions, is the time-resolved effective connectivity (EC) evoked by stimulating that target, using the Neural Perturbational Inference framework of Luo et al. [37] fit per participant. Right: the seed-based EC map evoked by stimulating one target (left temporo-parietal junction; red, positive; blue, negative response). **(B–D)** The plan of the study: *where*, *when* and *how* to stimulate for a reproducible effect. **(B)** *Where.* Group-mean EC matrix between cortical regions (ordered by Yeo network), and the per-target effect size on the cortex (mean GES, the total response a target evokes; Methods). **(C)** *When.* The same target evokes different responses depending on the brain’s instantaneous total activity (the baseline energy) just before the pulse. **(D)** *How.* Two strategies to reduce trial-to-trial variability: timing the pulse to low-energy states (closed-loop) and stimulating two regions at once (bifocal); the question marks denote the open question this work addresses.

To stimulate a region in silico, we add a fixed increment to its activity at a chosen time point *t* and propagate the input through the ANN. The difference between the perturbed and unperturbed states at *t* + 1 defines the predicted effect of stimulating that region at that time, which we read out as a seed-based effective connectivity (EC; Figure 1A, right).

Perturbing every region at every time point yields a complete map of the spatiotemporal dependence of stimulation effects, organized as one EC matrix per time point. An EC matrix (Figure 1B, top) captures the causal influence of each region on every other, as learned from the resting state data alone, and is not inflated by temporal autocorrelation.

We use this platform to ask three questions, laid out in Figure 1B–D. First, where in the cortex stimulation acts most strongly. For each target region, we analyze the topographic organization of its global effect size (GES), a scalar quantity that measures the magnitude of the evoked whole-brain response (Figure 1B, bottom). Second, when to stimulate to induce different responses. We ask whether the same perturbation evokes different effects depending on the brain’s instantaneous activity at the moment of stimulation (Figure 1C). Third, how the response can be made more reproducible. We explore two stimulation strategies: timing the pulse to the ongoing brain state and stimulating two regions together (Figure 1D).

### 2.1 The personalized models reproduce each brain’s resting-state dynamics

A personalized brain model is only useful if it behaves like the brain it was trained on. Starting from an initial state *t*, and recursively inserting the *t* + 1 output back into the ANN, each model generated resting-state BOLD timeseries resembling the empirical recordings (Figure 2A). The same framework was previously employed and validated on the same HCP dataset at the level of subject-averaged static FC [37]. We first confirmed that subject-averaged empirical and simulated FC matrices were correlated with *r* = 0.97. Next, we showed that the models accurately reproduced each participant’s static FC. For an example participant, the simulated FC matched both the pattern of the empirical FC (correlation between the two matrices *r* = 0.86; Figure 2B) and the distribution of its values (Kolmogorov–Smirnov distance KS = 0.11; Figure 2D). Across the cohort, the empirical and simulated FC matrices correlated at a median of *r* = 0.64 (Figure 2E), with a median KS distance of 0.22 (Figure 2F). This per-participant agreement is a stricter test than the group-averaged comparison reported previously.

**Figure 2:**
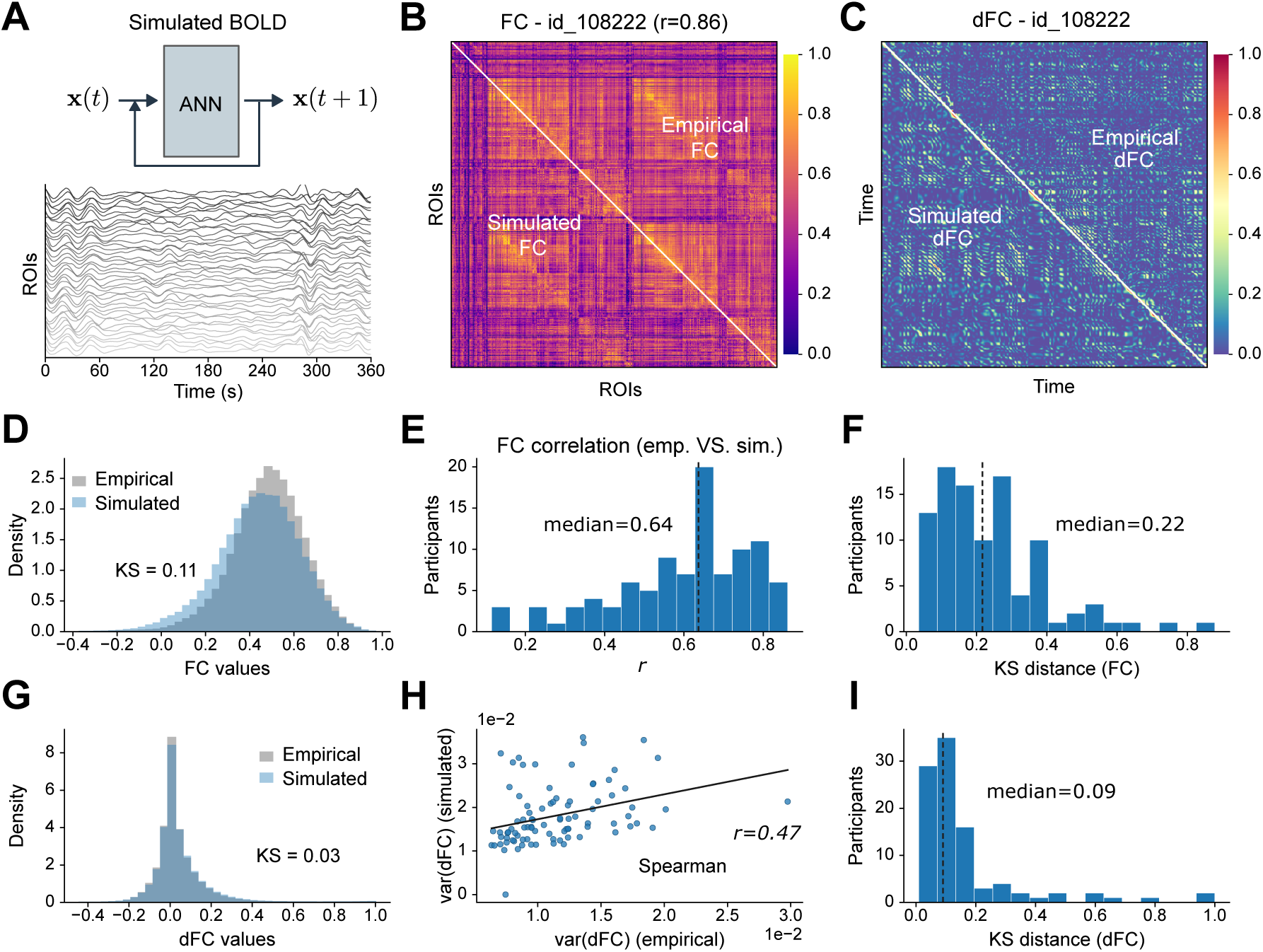
Personalized brain models reproduce each participant’s resting-state functional connectivity and its dynamics. **(A)** A trained model run in closed loop (each prediction fed back as the next input, with a small additive noise term) generates a surrogate BOLD time series (example participant; 360 s shown). **(B)** Static functional connectivity (FC) for an example participant: empirical (upper triangle) and simulated (lower triangle), Pearson correlations between regional time series; the two matrices correlate at *r* = 0.86. **(C)** Dynamic FC (dFC) for the same participant, empirical (upper triangle) versus simulated (lower triangle). **(D)** Distribution of FC values (all region pairs) for the example participant, empirical versus simulated; the distributions are close (Kolmogorov–Smirnov distance KS = 0.11). **(E)** Across the 100 participants, the correlation between each participant’s empirical and simulated FC (median *r* = 0.64; dashed line). **(F)** Across participants, the KS distance between empirical and simulated FC value distributions (median 0.22). **(G)** Distribution of dFC values for the example participant, empirical versus simulated (KS = 0.03). **(H)** Across participants, the variance of the empirical dFC against that of the simulated dFC, with linear fit (Spearman *r* = 0.47): the model reproduces how much the connectivity fluctuates over time. **(I)** Across participants, the KS distance between empirical and simulated dFC value distributions (median 0.09).

Since we are interested in the temporal dependence of stimulation, here we extended the validation at the level of subject-specific resting state network dynamics. Dynamic functional connectivity (dFC) measures how similar the brain’s connectivity patterns are between pairs of time points (Figure 2C). The distribution of dFC values was nearly identical between data and simulation (KS = 0.03 for the example participant, median 0.09 across the cohort; Figure 2G,I). The variance of the dFC, which quantifies how much the connectivity fluctuates and recurs over time, tracked the empirical values across participants (Spearman *r* = 0.47; Figure 2H).

The models were also individualized, with each model predicting its own participant’s held out activity better than that of any other participant and emerging as the best matching model for that participant, demonstrating that the framework captures individual rather than generic brain dynamics (Supplementary Figure S4).

### 2.2 Effective connectivity reproduces task-evoked activation across the cortex

To further validate the models against independent data, we asked whether virtually stimulating specific targets would induce seed-based EC maps that resemble the empirical task activation maps obtained from task fMRI in the same HCP participants across seven cognitive tasks (motor, working memory, language, relational, social, emotion, and gambling).

For each task condition, we defined an activation map over the 400 cortical regions following the four steps illustrated for the motor task in Figure 3A. Starting from the fixation task block design (step 1), we averaged the activity within each task block after accounting for a 5 s hemodynamic lag (step 2) and subtracted the fixation baseline (step 3). We then subtracted the average activation across the task’s conditions to isolate the activity specific to that condition (step 4). This procedure ensures that the resulting activation maps are distinguished from spontaneous fixation activity, while regressing out activity components shared across the task’s conditions.

**Figure 3:**
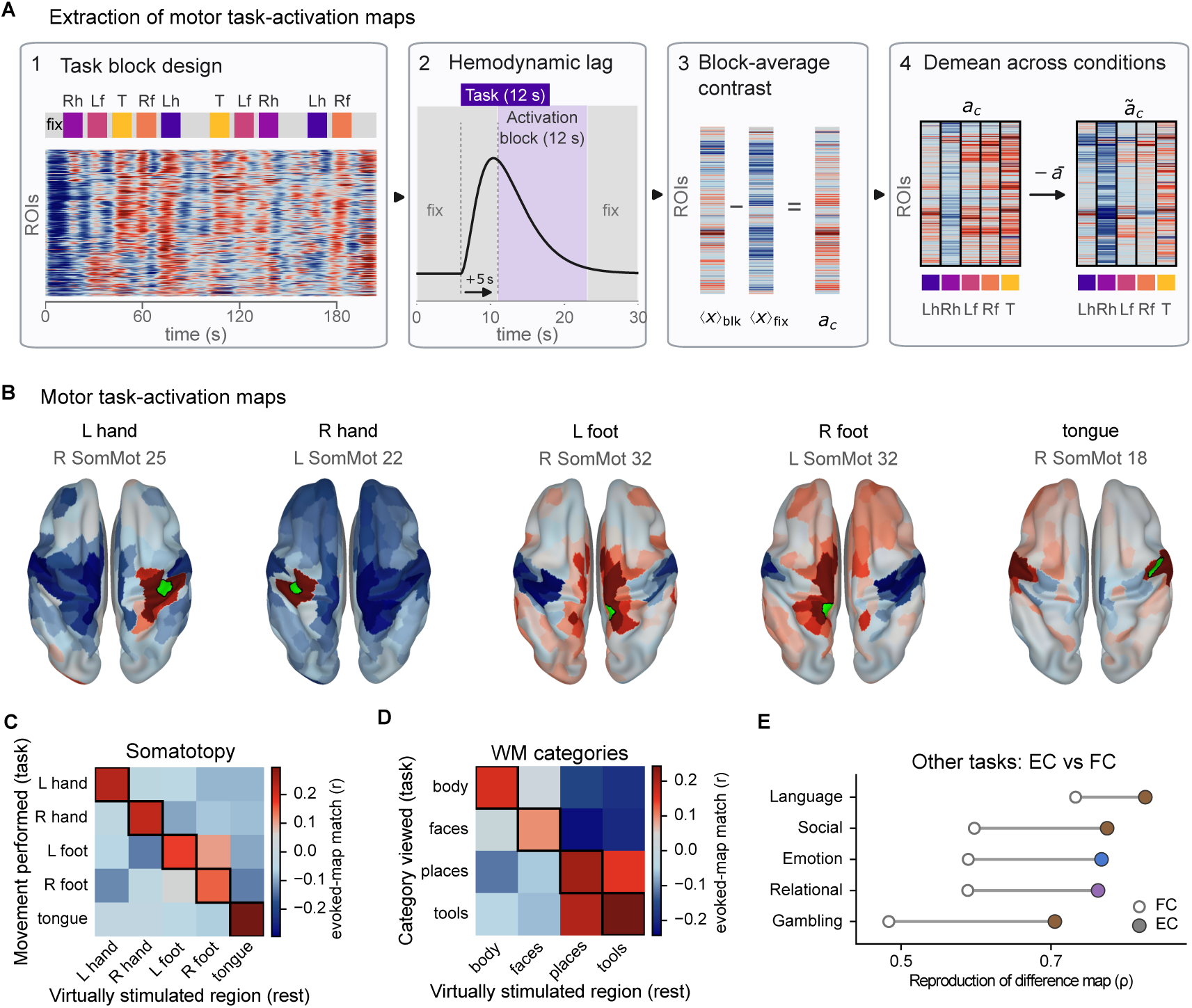
The models’ effective connectivity reproduces task-evoked activation and discriminates task conditions, more faithfully than functional connectivity. **(A)** Building the motor task-activation maps, in four steps. (1) The task is a block design alternating short blocks of each movement (left and right hand, left and right foot, tongue) with fixation. (2) Each block is read after a 5 s hemodynamic lag. (3) The activation map of a movement, *a_c_*, is its block-averaged activity minus the fixation average, one value per region. (4) Subtracting the across-movement mean *̄a* gives the demeaned map *̃a_c_*, isolating movement-specific activity. **(B)** Movement-specific activation maps for the five movements (dorsal view of both hemispheres; red, above the across-movement average; blue, below). Green marks the best stimulation seed for each movement—the region whose virtual-stimulation (EC) map most closely matches that movement’s activation—labelled with its Schaefer parcel. **(C)** Somatotopy. Confusion matrix of a five-way classification: each movement is assigned to the somatomotor seed whose virtual-stimulation map best matches its activation, with seeds fixed once on the group-average maps but each participant decoded from their own evoked and task maps (non-circular). Rows, movement performed; columns, region virtually stimulated; entries are the across-participant mean, the outlined diagonal marking correct assignment. Accuracy EC 91% versus FC 42% (chance 20%; FC matrices in Supplementary Figure S3); the only leakage is between the two feet, adjacent on the medial wall. **(D)** Working-memory categories, read as in (C) for the four picture categories (faces, places, bodies, tools); EC 68% versus FC 58% (chance 25%). **(E)** The five two-condition tasks. Across-participant median spatial similarity (Spearman correlation across the 400 cortical regions) between each task’s difference activation map and the stimulation-evoked EC map (filled) versus the resting FC map (open); EC exceeds FC for every task. Markers coloured by the task’s canonical Yeo-7 network.

From the model trained on resting state activity, we virtually stimulated each cortical region and extracted the resulting whole brain activity pattern, represented by the corresponding row of the seed based EC matrix. We then compared these stimulation evoked patterns with the task activation maps.

In the motor task, empirical activation maps elicited by movement of a given body part were recapitulated by virtual stimulation of the corresponding somatomotor region (Figure 3B). The five-way classification of the movements (left and right hand, left and right foot, tongue) via modeled EC reached 91% mean accuracy across participants (chance 20%), against only 42% with FC. Thus, the model recovers the motor homunculus, with the only confusion occurring between the two feet, which are adjacent on the medial wall (Figure 3C).

The same analysis applied to the working memory task, whose four picture categories (faces, places, bodies, and tools) engage distinct category selective regions, decoded the categories with an accuracy of 68% (chance 25%), compared with 58% using FC (Figure 3D).

The five remaining tasks contrasted two conditions: story versus math (language), win versus loss (gambling), fearful faces versus shapes (emotion), relational reasoning versus matching (relational), and social versus random motion (social). For each task, we defined a single difference activation map. In every case, the regions whose stimulation best reproduced the difference map were those most strongly activated by the task, and the EC map matched the activation more closely than the FC map in 100% of participants (Figure 3E). The best stimulation target also fell within the canonical network engaged by each task: frontoparietal for relational reasoning, default mode for language and social cognition, and visual for the emotion task, whose fearful faces versus shapes contrast prominently recruits visual and face processing regions (Supplementary Figure S1).

Having validated the models against resting state dynamics and task data, we next address the main questions of where, when, and how to stimulate to obtain the desired effects.

### 2.3 Where: the response increases along the cortical hierarchy

We first asked where stimulation acts most strongly. For each cortical target, we measured the GES averaged over stimulation times and across participants. The mean(GES) increased along the cortical hierarchy, from primary sensory and motor networks to transmodal association cortex (Figure 4A): lowest in the visual and somatomotor networks, intermediate in the attention and control networks, and highest in the limbic and default-mode networks [38]. The gradient was not a group-average artifact. Within individual participants, the rank correlation between a network’s position on the unimodal-to-transmodal axis and its responsiveness was strongly positive (median Spearman *ρ* = 0.82, positive in 97% of participants; Figure 4A, bottom).

**Figure 4:**
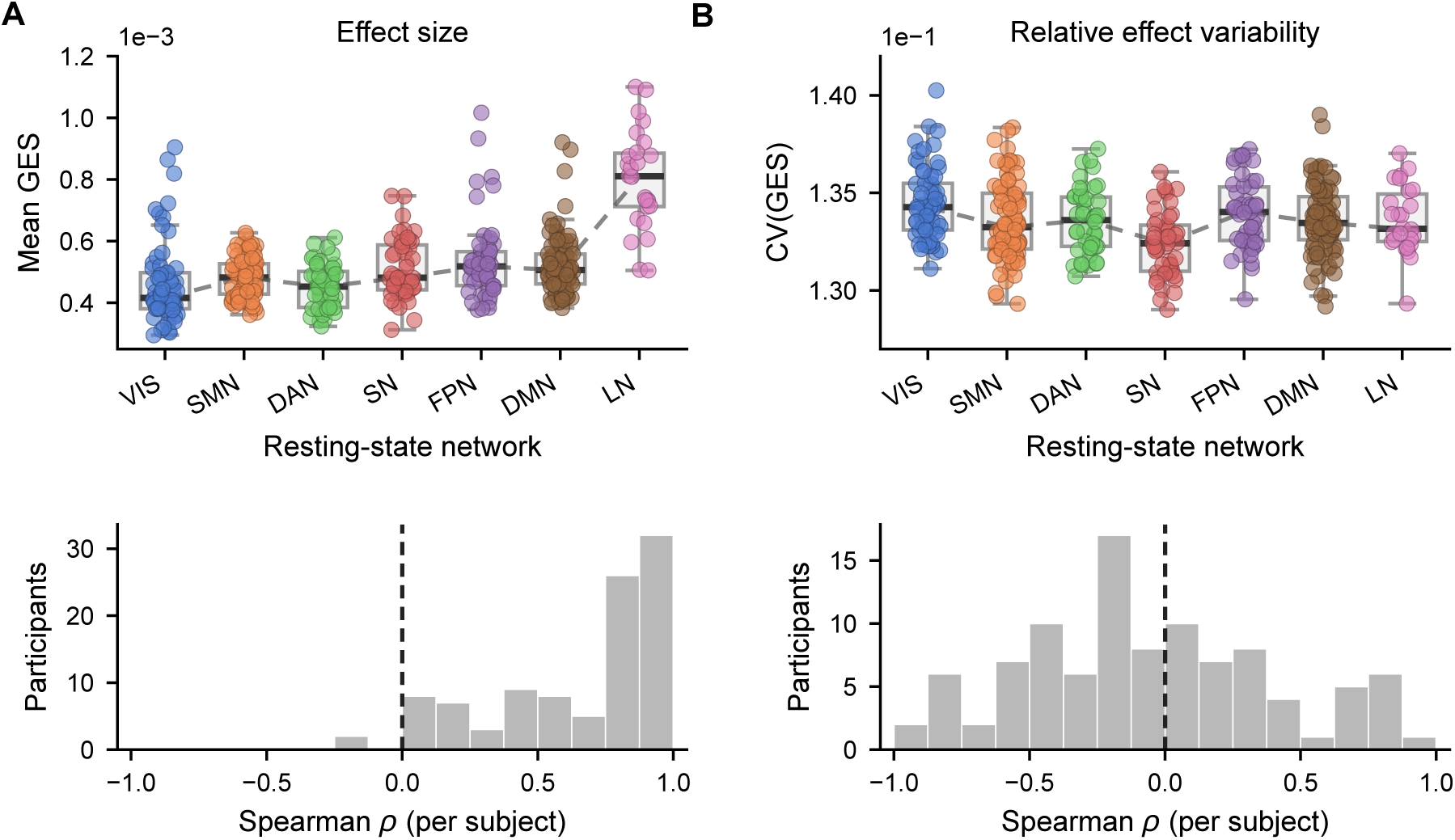
The magnitude of the response increases along the cortical hierarchy, while its relative variability is uniform. mean(GES) and CV(GES) are computed per participant over stimulation times and, where mapped per region, averaged across participants. **(A)** *Effect size.* Top: mean(GES) (responsiveness), one value per cortical region, grouped by Yeo-7 network and ordered from unimodal to transmodal (VIS, SMN, DAN, SN, FPN, DMN, LN); boxplots show median and interquartile range and the dashed line connects network medians. Bottom: per participant, the Spearman correlation between the seven network means and their hierarchy rank, well above zero (median *ρ* = 0.82, positive in 97%). **(B)** *Relative variability of the magnitude.* As in (A) for the coefficient of variation CV(GES) (standard deviation over mean of the effect size across stimulation times). The relative trial-to-trial variability of the response magnitude is uniform across networks (≈ 0.13) and has no per-participant hierarchy gradient (median *ρ* = −0.13, centred on zero).

By contrast, the *relative* trial-to-trial variability of the response, expressed as the coefficient of variation CV(GES) (i.e., the standard deviation of the effect size across stimulation times divided by its mean), was uniform across the cortex, sitting at ≈ 0.13 in every network, with no hierarchy gradient (median per-participant Spearman *ρ* = −0.13, centred on zero; Figure 4B). We notice that the absolute variance Var(GES) does climb the hierarchy, but only because it scales with the mean (Var(GES) ≈ (0.13 mean(GES))^2^); once this magnitude scaling is divided out no spatial structure remains. The response thus grows in size along the unimodal-to-transmodal axis, while the relative trial-to-trial variability of that size stays uniform across the cortex.

We also asked whether these regional properties track the cortical distribution of neuromodulator systems. We correlated mean(GES) and CV(GES) with 19 PET-derived receptor and transporter density maps, assessing significance against two spatial-autocorrelation-preserving nulls (a spin test and Moran spectral randomization), with FDR correction across maps (Supplementary Figure S6). The mean(GES) had no robust molecular correlate: its strongest, the dopamine transporter (DAT, *ρ* = +0.38), reached nominal significance under the spin null (*p*_spin_ = 0.029) but not the Moran null (*p*_Moran_ = 0.062) and did not survive FDR. Response reproducibility was more systematically organized: CV(GES) correlated negatively with several neuromodulator maps—most strongly the *µ*-opioid receptor (*ρ* = −0.29, *p*_spin_ = 0.001, *p*_Moran_ = 0.002, surviving FDR under both nulls), with H_3_ and mGluR_5_ also surviving FDR under the Moran null—so that receptor-richer regions tended to respond more reproducibly. This was a group-level spatial association only: it did not reproduce within individual participants (Supplementary Figure S6), so we treat these molecular associations as suggestive rather than definitive.

### 2.4 When: the brain’s ongoing activity gates the response

Having mapped where stimulation acts, we next addressed the when factor. We summarized the brain’s instantaneous state **x**(*t*) by its total ongoing activity just before the pulse, or its baseline energy 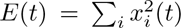,and related it to the size of the response. In a single participant and target, the GES fell steeply with baseline energy (Spearman *ρ* = −0.41; Figure 5A, left). The same relationship held when pooled across the cohort and all cortical targets (Figure 5A, middle), and in essentially every individual (per-participant correlation negative throughout the cohort; Figure 5A, right). In other words, the lower the ongoing activity, the larger the response. We refer to this dependence as gating: the brain’s ongoing state applies a continuous, state-dependent gain to the response, a whole-brain analogue of the gain modulation described in cortical circuits [39, 40].

**Figure 5:**
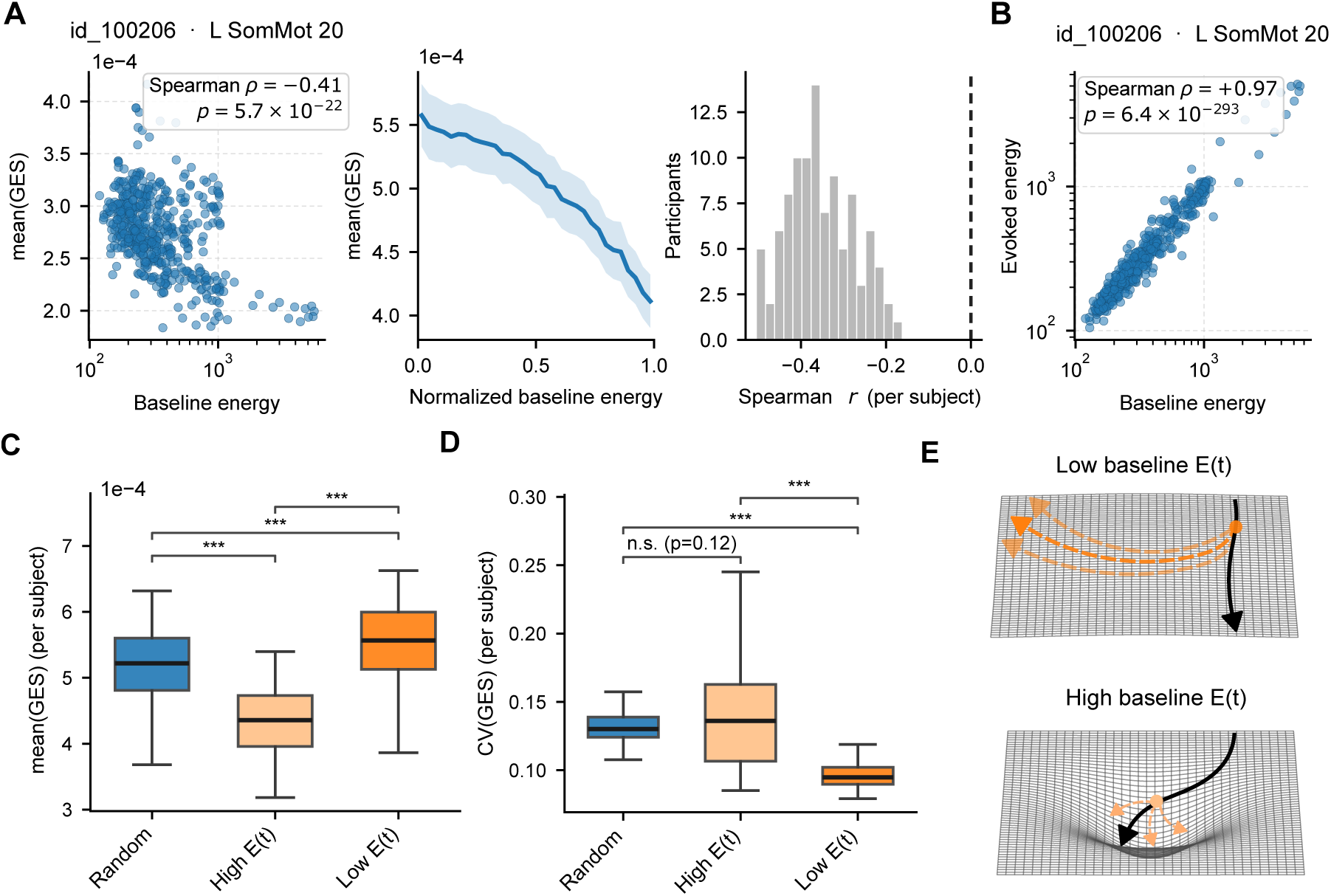
The brain’s ongoing activity gates the size and reproducibility of its response to stimulation. **(A)** The gating at three levels. Left: for one participant and target (L SomMot 20), the global effect size (GES) at each stimulation time against the pre-stimulation baseline energy (the brain’s total ongoing activity) on a logarithmic axis: the lower the ongoing activity, the larger the response (Spearman *ρ* = −0.41). Middle: the same relationship pooled across all participants and cortical targets (line, mean; band, spread across targets). Right: the per-participant baseline-energy–GES correlation lies entirely below zero, so the gating holds in essentially every individual. **(B)** For the same participant and target, the total evoked energy after stimulation tracks the baseline energy before it almost one-to-one (Spearman *ρ* = +0.97): a pulse does not reset the brain’s overall activity, so a given perturbation has a larger relative effect from a low-energy state. **(C)** Effect size by timing. For each participant, mean(GES) when stimulation is restricted to the 5% of time points with the lowest, the highest, or randomly chosen baseline energy; low-energy states give the largest effects and high-energy the smallest (boxplots, cohort; asterisks, pairwise Wilcoxon signed-rank). **(D)** Variability by timing, as in (C) for var(GES): low-energy stimulation is the least variable and random timing the most, all three comparisons significant. **(E)** Schematic on an energy landscape: from a low-energy state (top, flat) a perturbation produces a large but consistent displacement that stays low-energy, whereas from a high-energy state (bottom, steep) the same perturbation gives a smaller, more variable effect.

The gating reflected the configuration of the whole brain rather than the target’s own activity. Across cortical targets, the whole-brain baseline energy correlated with the response several-fold more than the target’s local activity (Supplementary Figure S7). This generalizes to whole-brain fMRI an effect reported in intracranial recordings, where pre-stimulus whole-brain features correlated with the response better than local ones [25].

Importantly, the gating is a nonlinear property of the learned dynamics, not an artifact of the perturbation procedure. A linear model (vector-autoregression) fit to the same data has, by construction, a perturbation response that does not depend on the ongoing state, so running the identical pipeline through it removes the gating entirely: the global effect size is constant across time for every target and participant, whereas in the model it falls with baseline energy in 99.8% of cortical targets (Supplementary Figure S8). The state-dependence therefore reflects nonlinear structure the model learns from the data. It is also not an artifact of the rarer high-energy states being poorly fit during the ANN training phase: the model predicts the dynamics equally well across the energy range (held-out cosine similarity 0.92 to 0.96 from low to high energy), and the gating persists when the analysis is restricted to the densely sampled central range of baseline energies (Supplementary Figure S9).

It should be noted that the total energy of the brain after stimulation tracked its energy before almost one-to-one (Spearman *ρ* = +0.97; Figure 5B): a pulse does not reset the brain’s overall activity. The same perturbation therefore produces a larger response relative to the ongoing activity when that activity is low, which is why low-energy states yield the largest effects.

The same models also let us ask whether the gating of the response generalizes beyond rest. Driving each model with the task states as initial conditions, the gating by ongoing activity and its unimodal-to-transmodal organization both persisted (Supplementary Figure S2B,C). The baseline energy that sets the gating itself tracked the cognitive state. It was lower during effortful, engaged conditions, falling by about half a standard deviation both when participants did mental arithmetic rather than listen to a story and during working-memory blocks relative to the fixation periods between them. It did not shift for brief movements, nor when working-memory load was raised from the 0-back to the 2-back condition (Supplementary Figure S2A). Because the response grows as baseline energy falls, the models predict that the same stimulation would evoke a larger and more reproducible effect during these engaged cognitive states than at rest, a directly testable prediction.

### 2.5 How: timing the pulse and pairing two regions

Because the response depends on the brain’s ongoing state, the timing of the pulse has the potential of reducing the variability of the brain’s response to stimulation. Restricting stimulation to the 5% of time points with the lowest baseline energy produced both the largest effect (Figure 5C) and the least variable one (lowest var(GES); Figure 5D), whereas random timing gave the most variable response and high-energy timing the smallest effect. Thus, timing the pulse to low-energy states, a closed-loop strategy, makes the response both larger and more reproducible (Figure 5E).

Closed-loop timing requires monitoring the brain state in real time, which is not always feasible in clinical or research settings. We therefore asked whether an open-loop alternative can also shape the response. In particular we focused on stimulating two regions at once (bifocal stimulation). The model let us evaluate the effect of stimulating every pair of cortical regions, about 80,000 in total, comparing the jointly evoked response with that of the single sites.

Bifocal stimulation changed both the size and the reproducibility of the response. Some pairs evoked a larger effect than the stronger of their two single sites (Figure 6A); For an example target in the left somatomotor cortex (SomMot LH 12), joint stimulation with its strongest-effect partner in the same network (SomMot LH 16) produced a substantially larger effect than stimulation of either site alone, whereas the two regions elicited similar effects when stimulated separately (Figure 6B). Moreover, in most target pairs, co-stimulation made the response more reproducible: the bifocal coefficient of variation, CV(GES), fell below that of the better single site for most stimulation pairs (Figure 6C). This is illustrated for an example target in the left somatomotor cortex and its optimal partner in the right lateral prefrontal cortex of the control network, where joint stimulation reduced CV(GES) below the values obtained with either site alone, despite the two single-site stimulations producing similar CV(GES) values (Figure 6D).

**Figure 6:**
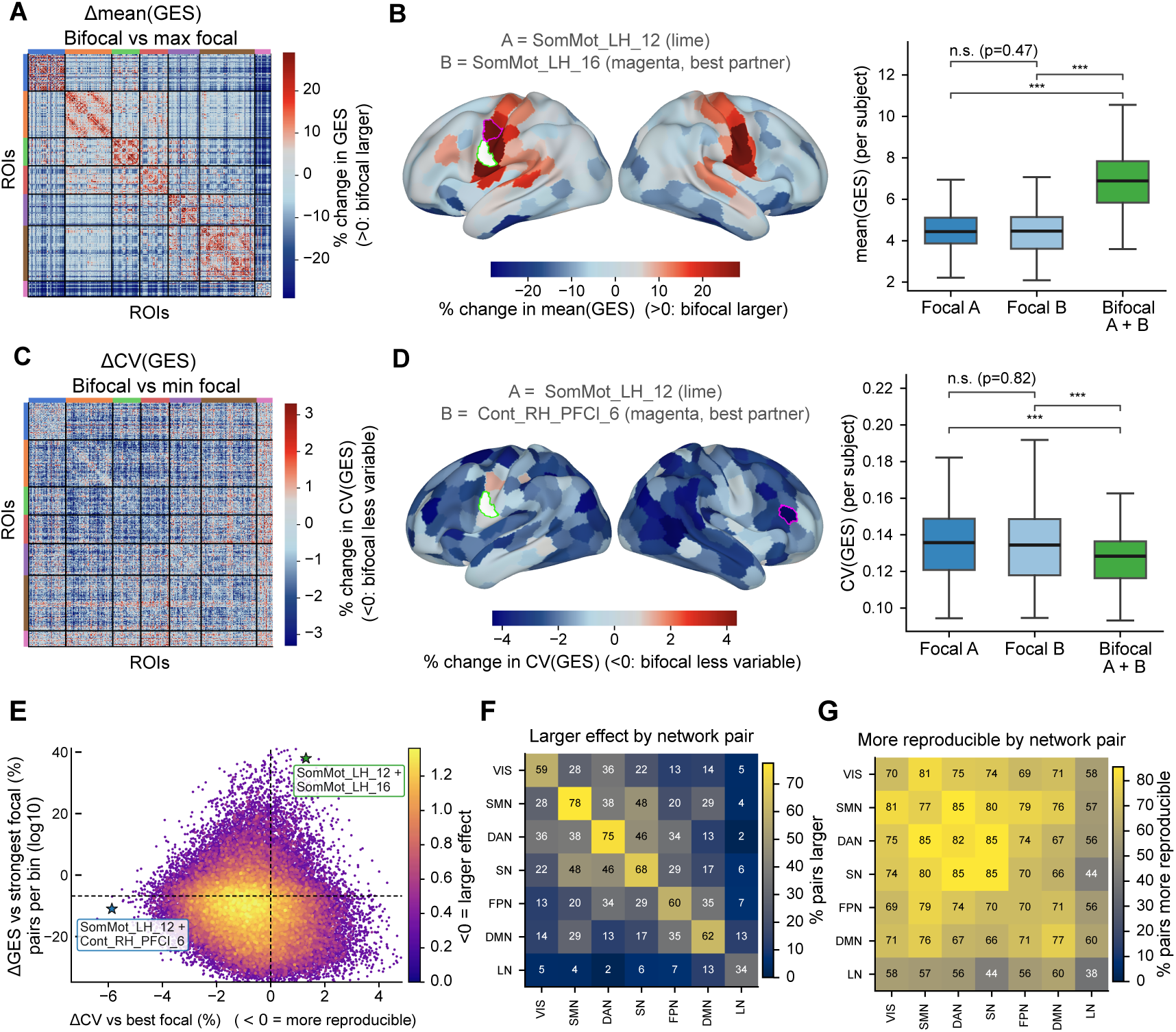
Bifocal stimulation amplifies the response within networks and makes it more reproducible across the cortex. Bifocal stimulation perturbs two targets at once; each pair’s jointly evoked response is compared with its single (“focal”) sites. The model evaluates all ∼80,000 cortical pairs, a search inaccessible to single-target experiments; matrices and surface maps are group averages. **(A)** Percentage change in mean(GES) of bifocal co-stimulation relative to the stronger single site, for every region pair (red, bifocal larger). **(B)** An example effect-increasing pair, A = SomMot LH 12 with its best effect partner B = SomMot LH 16. Left: percentage change in mean(GES) from pairing A (lime) with every other region (B outlined, magenta). Right: per-participant mean(GES) for A, B and A+B; the single sites do not differ whereas the pair is substantially larger (Wilcoxon signed-rank). **(C)** Percentage change in CV(GES) of bifocal relative to the more reproducible (lower-CV) single site (blue, bifocal more reproducible). **(D)** An example reproducibility-improving pair, A = SomMot LH 12 with its best CV partner B = Cont RH PFCl 6, as in (B) for CV(GES): pairing lowers CV(GES) below either single site. **(E)** The full landscape: for every pair, the change in CV relative to the better single site (horizontal; *<*0, more reproducible) against the change in mean(GES) relative to the stronger single site (vertical; *>*0, larger), coloured by pair density. The upper-left “win-win” quadrant holds 24% of pairs; stars mark the two example pairs. **(F)** For each pair of Yeo-7 networks, the fraction of region pairs whose bifocal effect exceeds the stronger single site; amplification concentrates within networks (diagonal, 62% within versus 20% across). **(G)** The fraction of region pairs whose bifocal CV(GES) is below the better single site, by network pair; reproducibility improves broadly (71–85% across most combinations), most among the sensory and attention networks.

Across all ∼80,000 pairs, the joint response was more reproducible than the better single site for 73% of pairs and larger than the stronger single site for 31%, and 24% fell in the win-win regime—jointly stronger *and* more reproducible (Figure 6E). For a given target the model therefore identifies many partner regions that make its effect both larger and more reliable.

Effect amplification was concentrated within networks: co-stimulating two regions of the same network exceeded the stronger single site far more often than cross-network pairs did (62% of within-network pairs versus 20% across networks; Figure 6F). Reproducibility gains, by contrast, were widespread: the bifocal CV fell below the better single site for a majority of pairs in almost every network combination (71–85%), most strongly among the sensory and attention networks (Figure 6G). Amplification is therefore network-specific while improved reproducibility is broadly distributed, and a large fraction of pairs achieve both.

## 3 Discussion

We used personalized brain models of resting-state fMRI to map how the brain’s response to stimulation depends on where and when it is delivered, and to test how it can be made more reliable. Three results stand out. The magnitude of the response increases along the cortical unimodal-to-transmodal hierarchy, while its relative trial-to-trial variability is uniform across the cortex. The response is gated by the brain’s ongoing state, whereby perturbations delivered when total activity is low evoke larger effects than identical perturbations delivered when activity is high, and this gating holds across participants and cortical targets. Finally, closed-loop and circuit-level stimulation approaches show promise for increasing effect size and the reproducibility of stimulation responses.

An important question is whether predictions from a framework built entirely in silico generalize to empirical brain dynamics. Several lines of evidence indicate that they do. Each personalized model reproduces its own participant’s resting functional connectivity and its dynamics (Figure 2). Beyond resting dynamics, the models also reproduce task-evoked activity. The EC estimated from virtual perturbations of models trained on resting state data reconstructs the activation patterns of seven cognitive tasks more faithfully than FC (Figure 3). The agreement is also anatomically specific. The best stimulation target for each task lies within the network it engages, and within the motor and working memory tasks the model discriminates individual conditions from the evoked maps, recovering the motor homunculus by classifying the five movements at 91% versus 42% for FC (chance 20%), and the four object categories at 68% versus 58% (chance 25%). Together, these findings indicate that virtual perturbations of models trained on resting state data recover the large-scale functional organization revealed by empirical task responses. This provides the strongest validation currently possible without a stimulation dataset and strengthens the interpretation of the model-derived predictions presented here. Future empirical validation will require testing these predictions with concurrent TMS-EEG, which provides a direct readout of cortical reactivity and effective connectivity, while also posing methodological challenges for data acquisition and analysis [41].

The surrogate-brain framework of Luo et al. [37] established that an ANN trained on fMRI can be used to estimate a brain’s EC, summarized as a single time-averaged matrix, which closely matches empirical structural connectivity. Here, we resolve the perturbation in time and condition it on the ongoing brain state, which turns the static connectivity matrix into a state-dependent quantity. Thus, the EC depends on the momentary brain state, rather than being a fixed property of the connectome. Fitting one model per participant, we established that the main reported effects, the cortical hierarchy and the state-gating, hold within individuals rather than on a group average.

The cortical hierarchy identified here is consistent with a broader organizing axis of the cortex. Functional connectivity varies along a principal gradient from unimodal sensory and motor regions to transmodal association cortex [38], and related gradients are present in cortical microstructure, myelination, gene expression, receptor density, and intrinsic timescales [42–45]. These anatomical and dynamical differences provide a plausible substrate for the graded stimulation effects observed in our models. Previous computational and perturbational studies similarly showed that the impact of local stimulation varies systematically with cortical hierarchy and regional timescale [46, 47]. Our results extend this work by showing that the magnitude of the immediate whole brain response increases from unimodal to transmodal targets within personalized models, while its relative variability remains approximately uniform across the cortex. The stimulation gradient may therefore reflect intrinsic differences in local circuit properties and large-scale embedding rather than differences in measurement reliability.

The state dependence of stimulation responses demonstrated here naturally suggests closed loop stimulation as a strategy for improving reproducibility. Restricting stimulation to low energy states made the response both larger and more reproducible, providing an in silico counterpart to empirical closed loop findings [14, 16]. Because real time monitoring of brain state is not always feasible, we also explored a complementary strategy based on circuit level bifocal stimulation. Stimulating two regions within a network amplified the response compared with focal stimulation of either region alone. Previous work has similarly moved from single site toward multi site stimulation [48]. For instance, multifocal tDCS of the motor network raises cortical excitability beyond conventional single site montages [49], and bilateral rTMS, delivering stimulation to two sites, has shown clinical benefit in treatment resistant depression [50]. Here we show that pairing regions within and across networks generally reduced response variability. More broadly, these findings suggest that a substantial fraction of response variability is not simply noise, but reflects a lawful dependence on the ongoing brain state and on the choice of stimulation targets, making it, in principle, controllable. This interpretation agrees with recent evidence that pre stimulation brain states predict and partly determine the variability of stimulation responses [3, 25]. Beyond response magnitude, evoked responsiveness covaries with non equilibrium features of ongoing dynamics, indicating that evoked responses and spontaneous dynamics reflect related aspects of brain state organization [51–53].

The use of brain models enabled exhaustive screening of a large number of initial conditions and stimulation targets, including the joint stimulation of all ∼80,000 region pairs, which is inaccessible experimentally [33]. Equally importantly, because the perturbed and unperturbed trajectories can be compared from the same initial state, the effect of each perturbation can be isolated directly without averaging across trials. This makes it possible to quantify the single-trial, state-dependent response that is inaccessible to conventional empirical analyses. Importantly, these predictions remain experimentally testable [3, 25]. The hierarchy and state gating of stimulation responses could be tested by contrasting stimulation with state-matched sham, or by using brain state-triggered closed-loop stimulation, and our results provide concrete predictions for such experiments.

Several aspects of the current implementation point to natural directions for future work. The individualized models were trained on the long resting state recordings available in the Human Connectome Project, which provide rich constraints on the learned dynamics. Such acquisitions, however, are uncommon in clinical settings, where substantially shorter scans may not constrain the models as well. Establishing how much data are required for a stable individualized fit will therefore be an important next step. The current implementation is also limited by the characteristics of the BOLD signal, which is slow, parcellated, and only indirectly related to neural activity [54, 55]. Consequently, the predictions concern the timescale of fMRI rather than the millisecond timescale at which neurostimulation operates. These limitations are not inherent to the framework itself. The same architecture can be trained on faster signals, and applying it to EEG, MEG, or intracranial electrophysiology would allow the same questions to be addressed at the timescale of the evoked responses themselves.

The virtual perturbation used here consists of a fixed transient increase in the BOLD activity of a single region. It is therefore a generic probe of the learned dynamics rather than a model of any particular stimulation device. Empirical stimulation acts on neural populations, engages both excitation and inhibition, and reaches the BOLD signal only through neurovascular coupling [56, 57]. Consistent with this, concurrent TMS-fMRI shows that a pulse does not reliably increase the BOLD signal at the stimulation site [58]. A natural next step is to tailor the perturbation to specific stimulation modalities by modifying its temporal profile, spatial extent, and amplitude while retaining the same personalized brain models.

A further limitation concerns how variability of the stimulus effects was quantified. Throughout, trial-to-trial variability refers to fluctuations in the magnitude of the response, summarized by the coefficient of variation of the global effect size. The variability of the evoked topography, namely how much the spatial response pattern changes from one brain state to the next, is a complementary measure that we did not examine here. Whether the spatial pattern of the response is as uniformly reproducible across the cortex as its magnitude therefore remains an open question.

Together, these results establish individualized surrogate brain models as a framework for systematically probing how the brain responds to stimulation. By revealing where, when, and how stimulation produces the most reliable responses, they provide a principled basis for designing future closed-loop and circuit-level stimulation protocols, while illustrating how personalized data-driven models can generate experimentally testable predictions about human brain dynamics.

## 4 Materials and Methods

### 4.1 Participants and data acquisition

We analyzed resting-state fMRI from 100 healthy adult participants of the Human Connectome Project (HCP) Young Adult release [59]. Data were acquired on a customized 3T scanner at 2 mm isotropic resolution with a repetition time of TR = 0.72 s. Each participant contributed four resting-state runs (two sessions, REST1 and REST2, each acquired with left–right and right–left phase encoding) of 1200 volumes each (≈14.4 min per run), for a total of 4800 volumes (≈ 57.6 min) of resting-state fMRI. Participants rested with their eyes open while fixating on a cross. For the validation of Figure 3, we additionally used the task fMRI of the same 100 participants across the seven HCP tasks (motor, working memory, language, relational, social, emotion and gambling); the task acquisition and the construction of the activation maps are described under *Reproduction of task-evoked activation* below. The models were trained only on the resting-state data.

### 4.2 Parcellation

Cortical regions were defined with the Schaefer-2018 atlas (400 parcels, organized into the seven Yeo resting-state networks) [60, 61]; subcortical regions with the Tian S3 atlas (50 parcels) [62], for *N* = 450 regions in total. The BOLD signal was averaged across the voxels of each parcel to give one representative time series per region. Each model was trained on all *N* = 450 regions, so subcortical activity contributes to the learned dynamics, but the perturbation analyses targeted and reported the 400 cortical Schaefer parcels only (in the full 450-parcel index, subcortical parcels are indexed 0–49 and cortical parcels 50–449). This mirrors the original NPI framework, where the subcortical regions are kept in training to minimize bias in the inferred cortical connectivity from unobserved areas, while the analysis focuses on the cortex [37].

### 4.3 Preprocessing

We started from the parcellated resting-state time series derived from the HCP minimally preprocessed, ICA-FIX-denoised data [59, 63]. For each run we discarded the first 30 volumes to remove non-steady-state effects, linearly detrended each regional time series, applied a zero-phase second-order Butterworth band-pass filter (0.008–0.08 Hz), and standardized each regional time series (z-score) within the run. The four preprocessed runs were then temporally concatenated, yielding a single time series of 4,680 volumes (≈56 min) per participant. No global-signal regression was applied.

### 4.4 Personalized neural-network model

For each participant we trained one feedforward ANN (a multilayer perceptron, MLP) to predict the brain’s activity one step ahead from its recent history. The input concatenates the *S* = 3 most recent activity vectors **x**(*t*), **x**(*t* − 1), **x**(*t* − 2) ∈ R*^N^* (*N* regions per frame), and the output is the next activity vector **x**(*t*+1) ∈ R*^N^* ; for clarity we refer to the most recent input frame as the current state **x**(*t*). The network has two hidden layers (widths 2*N* = 900 and 0.8*N* = 360) with rectified-linear (ReLU) nonlinearities and a linear read-out to the *N* = 450 outputs. We trained it to minimize the mean-squared one-step prediction error using the Adam optimizer (learning rate 5 × 10^−4^, *L*_2_ weight decay 5 × 10^−5^, batch size 64, 50 epochs) on an 80/20 split of the time points into training and held-out test sets. The trained network is the participant’s personalized brain model.

We adopted this architecture and training procedure directly from the Neural Perturbational Inference (NPI) framework of Luo et al. [37], who implemented the surrogate brain as an MLP. On the same HCP dataset they compared the MLP against convolutional, recurrent and vector-autoregressive surrogate models on signal prediction, functional-connectivity reproduction and effective-connectivity inference. While the NPI framework proved robust across all of these architectures, the MLP performed best on functional-connectivity reproduction and effective-connectivity inference, and was therefore chosen as the surrogate model. We adopt the same choice here. On synthetic data with known ground-truth effective connectivity, NPI matched the ground truth (correlation *r* = 0.95) and outperformed Granger causality and dynamic causal modelling, and its inferred connectivity reproduced cortico-cortical evoked-potential propagation [37]. Because we use the same data and the same model class, this validation transfers to our models, and we keep the architecture fixed rather than re-optimizing it for each participant.

### 4.5 Simulating resting-state activity

To generate surrogate resting-state activity, each trained model was run in closed loop (Figure 2A): starting from an initial state **x**(*t*), the predicted next state **x**(*t*+1) was fed back as the next input, with a small Gaussian noise term (standard deviation 0.1; since each regional time series was z-scored to unit variance, this is a tenth of every region’s standard deviation) added at each step to sustain the dynamics. Iterating this map produces a simulated BOLD time series of arbitrary length, generated entirely by the ANN without further input from the data.

### 4.6 Functional and dynamic functional connectivity

We initially validated each model by comparing the functional connectivity (FC) and dynamic functional connectivity (dFC) of its simulated activity with those of the participant’s empirical data (Figure 2B,C). Static FC was the *N* ×*N* matrix of Pearson correlations between regional time series. For each participant, we correlated the upper-triangular entries of the empirical and simulated FC matrices, yielding a single goodness-of-fit value per participant, and reported the distribution of this correlation across the cohort. As an additional check, and for direct comparison with [37], we also correlated the cohort-averaged empirical and simulated FC matrices, recovering the high group-level agreement reported there (*r* = 0.97).

Dynamic FC was computed with an edge-centric, window-free approach [64–66]. For each pair of regions *i, j* we define an edge co-fluctuation time series as the element-wise product of their standardized signals, **c**^(*t*) = *c_ij_*(*t*) = *x_i_*(*t*) · *x_j_*(*t*); at each time point, the vector of all edge cofluctuations (of size *N* × (*N* − 1)*/*2) is an instantaneous brain co-activation pattern, which we refer to as the instantaneous network state. The dFC is the matrix of Pearson correlations dFC(*t_a_, t_b_*) = *r*(**c**^(*t_a_*), **c**^(*t_b_*)) between these network states taken at every pair of time points, so that entry (*t_a_, t_b_*) measures how similar the brain’s configuration is at the two moments, without any sliding-window averaging. We summarized each dFC matrix by the variance of its off-diagonal entries, a measure of the amount of dynamic fluctuation, and compared empirical and simulated values across participants (Figure 2C). We screened each model for free-running stability (Tukey outliers in the variance of the simulated dFC); all 100 participants analysed here passed this screening.

### 4.7 Virtual perturbation and effective connectivity

A virtual perturbation at target region *j* adds a fixed increment (amplitude 0.1, in units of the standardized signal) to region *j* of the current state **x**(*t*) and propagates the input one step through the model to obtain the perturbed prediction **x**^(*j*)^(*t*+1). The unperturbed prediction **x**(*t*+1) is obtained from the same input without the increment. The effect of stimulating seed *j* on a downstream region *i* at time *t* is the time-resolved effective connectivity

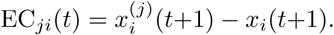

Evaluating this for every seed *j*, downstream region *i* and time *t* yields the time-resolved effective-connectivity tensor; its average over time (here we use 500 time points corresponding to different initial conditions) gives the static EC matrix (Figure 1B). Bifocal effective connectivity is defined by perturbing two seeds *j*_1_ and *j*_2_ simultaneously and evaluating the modulus of the effect (perturbed minus unperturbed) vector

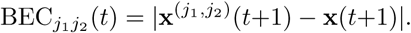

All perturbation analyses used the 400 cortical targets. We verified that results are stable across perturbation amplitudes (Supplementary Figure S5).

### 4.8 Symbols and response measures

The symbols used throughout the perturbation analysis, the local and global energy and effect-size measures derived from each perturbation, and the per-target summary statistics (including the variance-explained measures compared in Supplementary Figure S7) are collected in Supplementary Table S1. The quantities used in the main text are also defined where they first appear.

### 4.9 Global effect size and its summary statistics

The instantaneous global effect of stimulating target *j* at time *t* is the global effect size (GES) 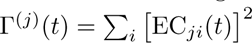, the total squared response evoked across the brain. For each participant and target we summarized the GES over stimulation times by its mean, mean(GES) = ⟨Γ^(*j*)^⟩*_t_*, which we use as the region’s *responsiveness*, and by two measures of its trial-to-trial variability: the absolute variance 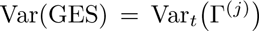, and the coefficient of variation 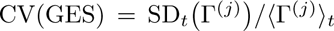, its *relative* variability. Because CV(GES) is nearly constant across regions, Var(GES) ≈ (CV · mean(GES))^2^ scales with the magnitude; we therefore use CV(GES) where variability must be separated from response size (the regional organization and receptor associations, Figure 4B and Supplementary Figure S6) and Var(GES) for the magnitude of induced variability and its reduction by the interventions (Figures 5 and 6).

### 4.10 State-dependence of the response

We summarized the brain’s instantaneous state immediately before each perturbation by the base-line energy 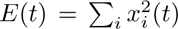, the total activity across regions. To quantify gating, we pooled all (participant, cortical target, time) samples and related the pre-stimulation baseline energy *E*(*t*) to the resulting global effect size Γ^(*j*)^(*t*+1), summarizing the relationship both by binning into quartiles of *E*(*t*) and by the within-subject and within-target rank correlations between the two quantities. To test whether the gating reflects the whole-brain state or the target’s own activity, we compared, for each target, the variance in the response explained by the global baseline energy *E*(*t*) with that explained by the target’s local baseline energy 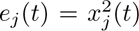. To map the gating onto cortical organization, regions were ordered along the principal functional gradient [38] and grouped by Yeo-7 network [61].

To test whether the gating is a nonlinear effect rather than an artifact of the perturbation pipeline, we fit an optimal linear one-step model to each participant (ordinary least squares with the same *S* = 3 window and intercept) and applied the identical virtual perturbation and GES computation. For a linear map the evoked response EC = *A **δ*** is independent of the current state, so Γ^(*j*)^(*t*) is constant across time; we confirmed this for all 100 participants (temporal coefficient of variation of the GES below 10^−13^ at every cortical target). For the MLP we summarized the gating per cortical target as the Spearman correlation between *E*(*t*) and Γ^(*j*)^(*t*) (Supplementary Figure S8). We further checked that the gating is not produced by the rarer high-energy states being poorly fit (Supplementary Figure S9): we recomputed the gating correlation restricted to the central 80% and central 60% of each participant’s baseline-energy distribution, and we evaluated the held-out one-step prediction accuracy of each model by baseline-energy decile, using the scale-invariant cosine similarity between predicted and observed next-state vectors (which, unlike the squared error, is not inflated by the larger signal magnitude of high-energy states).

### 4.11 Variability-reduction strategies

We evaluated two in-silico strategies for reducing the trial-to-trial variability of the induced effect, quantified by Var(GES). For *closed-loop timing*, we compared three sets of stimulation trials, each comprising 5% of the time points: those with the lowest baseline energy (low-energy), those with the highest (high-energy), and trials drawn at random (open-loop, averaged over repeated draws). For each (participant, target) we computed mean(GES) and var(GES) within each set, averaged them across cortical targets per participant, and compared the three regimes across participants with pairwise Wilcoxon signed-rank tests (Figure 5C,D). For *bifocal stimulation*, we used the bifocal effective connectivity BEC*_j_*_1_ *_j_*_2_ (*t*) to evaluate the joint stimulation of region pairs, and, for each cortical network, identified pairs whose joint response had a magnitude comparable to single-site stimulation but a lower Var(GES).

### 4.12 Receptor map associations

We tested whether two node-level properties of the response, its mean magnitude (mean(GES) = ⟨Γ^(*j*)^⟩) and its relative variability (CV(GES)), track the cortical distribution of neurotransmitter systems, using a group-level atlas of 19 PET-derived receptor and transporter density maps [67, 68] parcellated to the 400 Schaefer cortical regions. For each map and each measure we computed the Spearman correlation across the 400 cortical regions, using the across-participant mean of the measure.

Significance was assessed against two spatial-autocorrelation-preserving nulls—the Alexander–Bloch spin test and Moran spectral randomization (see *Statistics*)—and corrected across the 19 maps with the Benjamini–Hochberg false-discovery rate. These group-level associations are shown, for all 19 maps, in Supplementary Figure S6.

As a complementary, participant-resolved test, we also correlated each receptor map with the regional measure *within* each participant, obtaining one Spearman correlation per participant, and tested whether this distribution differed from zero across the cohort with a Wilcoxon signed-rank test, again FDR-corrected across the 19 maps. Both levels of analysis, for both the mean effect size and its relative variability, are shown in Supplementary Figure S6.

### 4.13 Reproduction of task-evoked activation

We used the task fMRI of the same 100 participants across the seven HCP tasks [69] (LR run; identical Schaefer-400/Tian-S3 parcellation and preprocessing to the resting data). Each task is a block design in which the participant performs one condition per block. In the *motor* task a visual cue instructs one of five movements (tapping the left or right fingers, squeezing the left or right toes, or moving the tongue); *working memory* is an *N* -back task on pictures from four categories (faces, places, body parts, tools) at two memory loads (0- and 2-back); *language* alternates listening to a short story with solving a spoken arithmetic problem; *emotion* matches fearful or angry faces versus neutral shapes; *relational* contrasts relational reasoning with a perceptual matching control; *gambling* contrasts monetary reward (win) with loss; and *social* contrasts clips of shapes that interact (theory of mind) with shapes moving randomly. Each task run is 176–405 volumes (≈ 2–5 min), an order of magnitude shorter than the 4,680-volume resting series used for training, so no model was fit to the task data; the models serve only as fixed predictors. The activation map of a condition is built in the four steps illustrated for the motor task in Figure 3A: the parcellated activity is averaged over that condition’s block volumes (each shifted by a 5 s hemodynamic delay) and referenced to the fixation baseline, and the mean across that task’s conditions is then subtracted. Subtracting this common mean removes the component shared by every block of the task—the generic engagement, arousal and broadly distributed default-mode and attention responses that any condition evokes—and leaves the activity specific to each condition; for a two-condition task the operation reduces to the standard contrast between the two (for example story minus math, the “positive” minus the “negative” condition). This subtraction matters because the shared component is large and falls on broadly connected transmodal hub regions, whose connectivity correlates with almost any activation map; regressing it out lets the comparison reflect the condition-specific network rather than these hubs.

#### Two-condition tasks

For each seed *j* we measured the spatial similarity 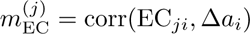 between its virtual perturbation effect (i.e., the row j of the EC matrix) and the task’s difference map 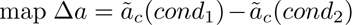. The same computation with FC gave *m*_FC_ (Figure 3E). We report the across-participant median for EC and FC and the fraction with EC*>*FC; the reproduction was also significant against an Alexander–Bloch / Vázquez-Rodŕıguez spin null (1000 rotations of the parcel centroids). The single best stimulation target per task, identified after discounting broadly connected hubs, is mapped per region in Supplementary Figure S1.

#### Within-task classification (motor, working memory)

For the tasks with several distinct conditions we asked whether the model can identify which condition is active from the region that best reproduces it—a nearest-template classification. From the group we fixed one stimulation seed per condition: the cortical region whose group EC evoked map correlated most with that condition’s group activation (restricted to somatomotor cortex for the motor movements, where the seeds are the body-part representations). Holding these seeds fixed, each participant was classified independently and without circularity: for every condition we correlated that participant’s condition activation with each seed’s evoked map and assigned the condition to the best-matching seed. The fraction correctly assigned—the diagonal of the confusion matrix—is the accuracy; we report its across-participant mean against the chance level (one over the number of conditions, 20% for motor and 25% for working memory) and a label-permutation null. Repeating the identical classification with each participant’s resting FC map in place of the EC map, and with seeds fixed from the group FC, quantifies the gain of effective over functional connectivity (Figure 3C,D; the FC confusion matrices are in Supplementary Figure S3).

### 4.14 Cognitive-state modulation of the gating

To test whether the gating generalizes beyond rest and tracks cognitively meaningful states, we drove each rest-trained model with the task states of the two multi-block tasks (language and working memory) and repeated the gating analysis. Task windows were built exactly as for rest—the *S* = 3 frames ending at each within-block volume, lagged by 5 s—and for each window we computed the baseline energy *E*(*t*) and the global effect size at all 400 cortical targets. We summarized the gating per participant as the Spearman correlation *ρ*(*E*(*t*), GES) across all task states (GES averaged over targets), and the hierarchy as the Spearman correlation between the seven Yeo-network mean responsiveness values (from the during-block states) and their unimodal-to-transmodal rank (Supplementary Figure S2B,C). Separately, and without the model, we tested whether the baseline energy itself differs across cognitive states (Supplementary Figure S2A): within each participant we contrasted the mean *E*(*t*) between conditions—math versus story (language); the N-back blocks versus fixation and 2-back versus 0-back (working memory); and movement versus fixation (motor)—expressing each difference in units of that participant’s temporal standard deviation of *E* and testing it across the cohort with a Wilcoxon signed-rank test. For the task-versus-fixation contrasts, fixation was restricted to interior inter-block volumes (between the first and last block), excluding run-edge non-steady-state volumes that otherwise inflate the resting energy; the within-task math-versus-story contrast is free of this confound by construction.

### 4.15 Statistics

Associations between regional maps are reported as Spearman rank correlations across the 400 cortical regions, except for the geometric-mean scaling of the bifocal responses (Figure 6E,F), which uses Pearson correlation. Because cortical maps are spatially autocorrelated, naive correlation *p*-values are anticonservative, so the significance of every spatial-map association (the receptor-density correlations and the task-activation reproduction) was assessed against a spatial-autocorrelation-preserving null. The primary null is the spin test: parcel centroids were projected onto the fsLR-32k spherical surface and randomly rotated 1000 times (Alexander–Bloch / Vázquez-Rodŕıguez [70]), and the two-tailed *p*_spin_ is the fraction of rotations whose absolute correlation equals or exceeds the observed one. For the receptor-density correlations we additionally report a Moran spectral-randomization null (2000 surrogate maps per receptor, generated from the PET density maps with the neuromaps implementation [68, 71]), with *p*_Moran_ defined analogously; a receptor association is called robust only if it survives FDR under this null. When a family of tests was run across the 19 receptor maps, *p*-values were corrected with the Benjamini–Hochberg false-discovery rate (*q*_FDR_) [72]. Paired, per-participant quantities were compared across the cohort with two-sided Wilcoxon signed-rank tests: the three closed-loop timing regimes (pairwise; Figure 5C,D), the bifocal example pairs (Figure 6B,D), the per-participant receptor correlations against zero (Supplementary Figure S6), the cognitive-state energy contrasts against zero (Supplementary Figure S2A), and the subject-specificity comparison (Supplementary Figure S4). Within-task classification accuracies were tested against the chance level (one over the number of conditions) and a label-permutation null. The significance threshold is *p <* 0.05 throughout; in the figures, asterisks mark ^∗^*p <* 0.05, ^∗∗^*p <* 0.01 and ^∗∗∗^*p <* 10^−3^, and n.s. marks *p >* 0.05.

## Acknowledgments

This work was supported by the Marie Skłódowska–Curie Postdoctoral Fellowship (Project CAERUS) under the European Union’s Horizon Europe research and innovation programme, grant agreement No. 101199894. I.A.-P. was supported by grant PID2022-136216NB-100 funded by MI-CIU/AEI/10.13039/501100011033 and by the European Regional Development Fund (ERDF) “A way of making Europe” (EU). G.D. was supported by grant PID2022-136216NB-I00 funded by MI-CIU/AEI/10.13039/501100011033 and by the European Regional Development Fund (ERDF) “A way of making Europe” (EU); by the ERC Synergy grant NEMESIS (ref. 101071900) funded by the European Union (Horizon Europe); and by the AGAUR research support grant 2021 SGR 00917 funded by the Department of Research and Universities of the Generalitat of Catalunya. Data were provided by the Human Connectome Project, WU–Minn Consortium (Principal Investigators: David Van Essen and Kamil Ugurbil; 1U54MH091657) funded by the 16 NIH institutes and centers that support the NIH Blueprint for Neuroscience Research, and by the McDonnell Center for Systems Neuroscience at Washington University.

## Author Contributions

G.R. conceived and designed the study. G.R., I.A.-P. and T.B.-B. performed the analyses and prepared the figures. G.R. wrote the original draft. G.D. supervised the work. G.R. and G.D. acquired funding. All authors reviewed and edited the manuscript.

## Data Availability

The resting-state fMRI data analysed in this study are available from the Human Connectome Project (WU–Minn HCP 1200 Subjects Release) at https://db.humanconnectome.org under the HCP Open Access Data Use Terms. The derived data supporting the findings (parcellated time series, trained per-participant models, and effective-connectivity matrices) are available at the code repository below and from the corresponding author on reasonable request.

## Code Availability

The full code to reproduce the data analysis described in this paper will be made freely and publicly available in an online repository upon acceptance of the manuscript.

## Supplementary Materials

Supplementary Table S1. lists the symbols and response measures used in the perturbation analysis. The supplementary figures report the controls and robustness analyses referenced in the main text: the per-region task-activation reproduction maps for all seven tasks (Figure S1) and the modulation of the gating axis by cognitive state, with the gating and hierarchy persisting for task-driven states (Figure S2), and the functional-connectivity counterpart of the within-task decoding (Figure S3); subject specificity of the models (Figure S4), stability of the perturbation analysis across amplitudes (Figure S5), the neurotransmitter-receptor associations (Figure S6), the whole-brain versus local origin of the gating (Figure S7), the linear-model and sampling controls for the gating (Figures S8 and S9), and the cohort-level statistics of bifocal stimulation (Figure S10).

**Figure S1:**
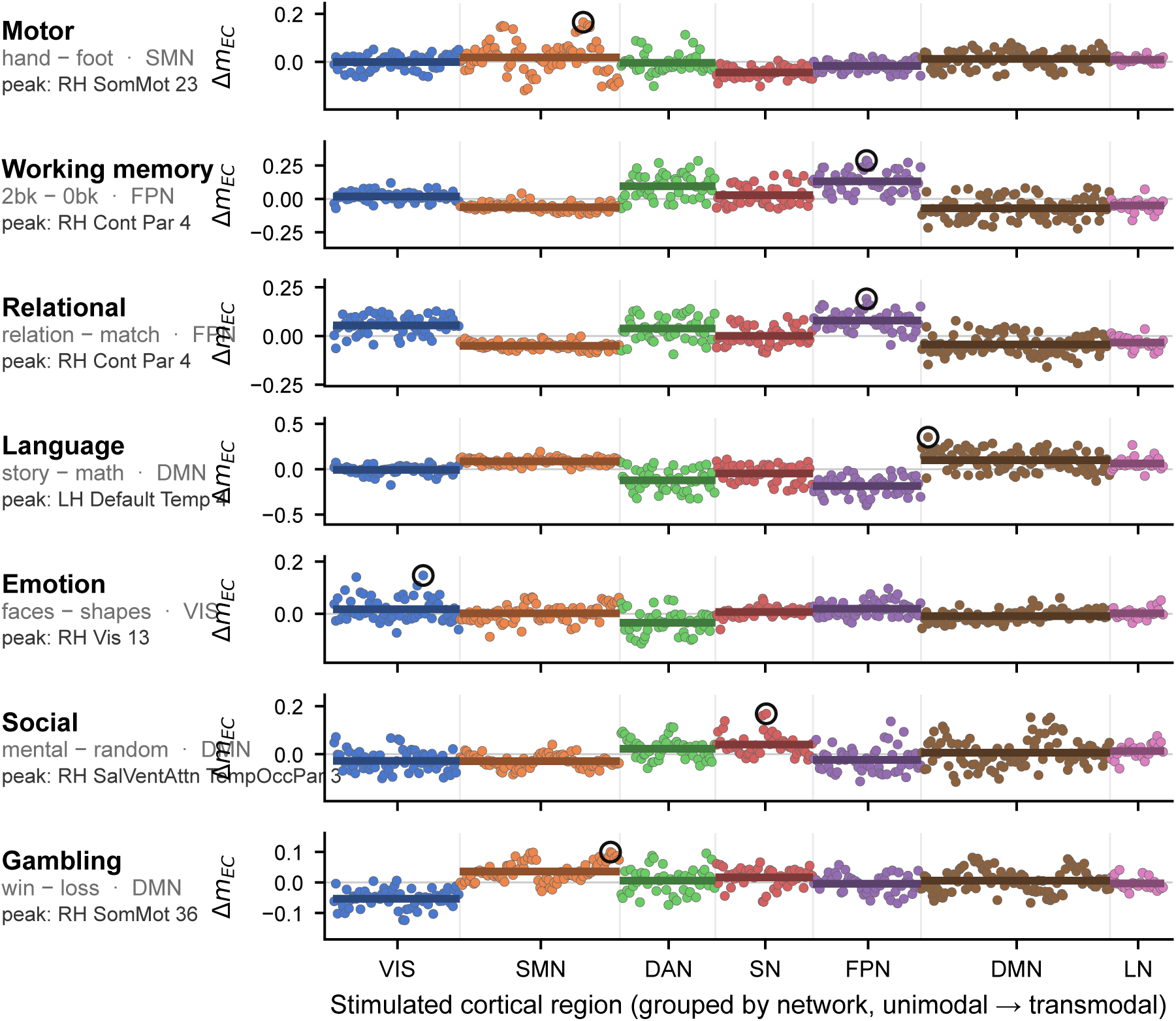
Per-region maps of task-activation reproduction, all seven tasks. For each task, the task-specific score Δ*m*_EC_ (the evoked-map similarity of each region to the task activation, minus that region’s mean across tasks) for every cortical region, grouped by Yeo-7 network (unimodal to transmodal) and colored by network. Dark horizontal lines are network means; the circled point is the peak region (labeled). The canonical network is elevated for the strong contrasts (motor, working memory, language, relational) and weaker for the subtler event-related contrasts (social, gambling).

**Figure S2:**
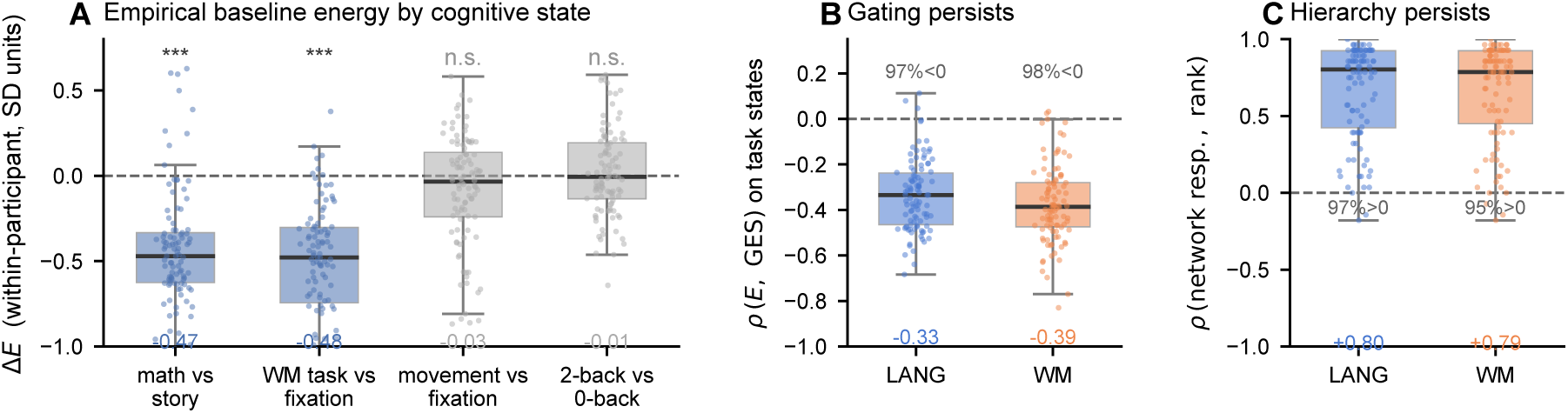
Cognitive state modulates the gating axis. **(A)** *Model-free.* Within-participant change in the empirical baseline energy *E*(*t*) =Σ*_i_x_i_*(*t*)^2^ between cognitive states, in units of each participant’s temporal standard deviation of *E* (box, cohort median and interquartile range; points, the 100 participants; dashed line, no change). Engagement and demand lower the baseline energy—arithmetic below narrative listening (math vs story, median −0.47 SD) and working-memory blocks below interior inter-block fixation (WM task vs fixation, −0.48 SD)—whereas brief movements (movement vs fixation) and parametric working-memory load (2-back vs 0-back) do not (medians printed below each box; ^∗∗∗^*p <* 10^−3^, n.s. *p >* 0.05, Wilcoxon signed-rank versus zero). Fixation is restricted to interior inter-block volumes to exclude run-edge non-steady-state states (Methods). Because the response grows as baseline energy falls, these lower-energy engaged states are the ones predicted to be more stimulable. **(B)** *The gating law generalizes to task-driven states.* Feeding each rest-trained model the task states and computing the global effect size (GES) at every cortical target, the per-participant Spearman correlation *ρ*(*E*(*t*), GES) remains negative for both the language and working-memory tasks (medians −0.33 and −0.39; negative in 97% and 98% of participants), as at rest. **(C)** *The hierarchy persists.* The per-participant Spearman correlation between the seven network-mean responsiveness values and their unimodal-to-transmodal rank stays positive on task states (medians +0.80 and +0.79; positive in 97% and 95%), matching the resting hierarchy.

**Figure S3:**
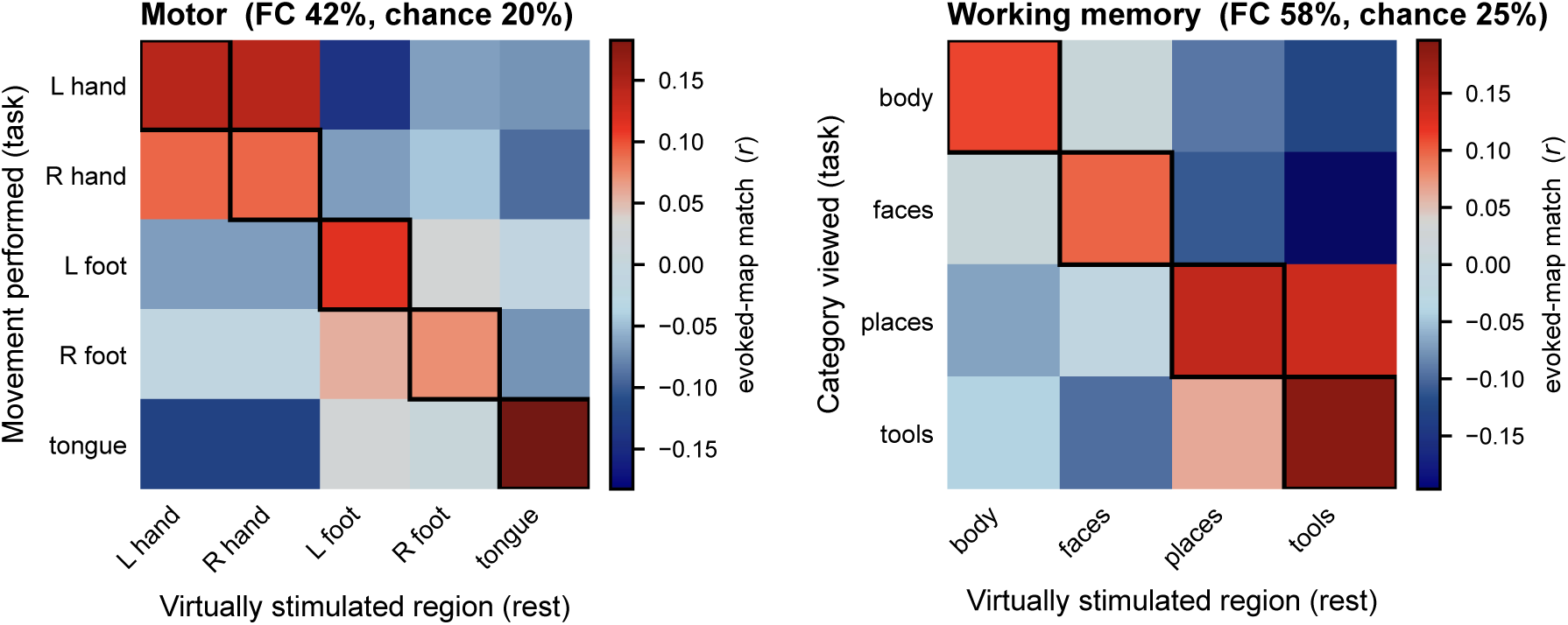
Within-task decoding from functional connectivity (the FC counterpart of Figure 3C,D). Confusion matrices for the motor (left) and working-memory (right) decoding when each condition’s seed is the resting functional-connectivity map rather than the effective-connectivity map. Functional connectivity, lacking causal direction, smears the two hands and the two feet together (motor accuracy 42%, against 91% for effective connectivity) and separates the working-memory categories only coarsely (58% versus 68%). Rows, condition; columns, stimulated region; outlined diagonal, correct assignment.

**Figure S4:**
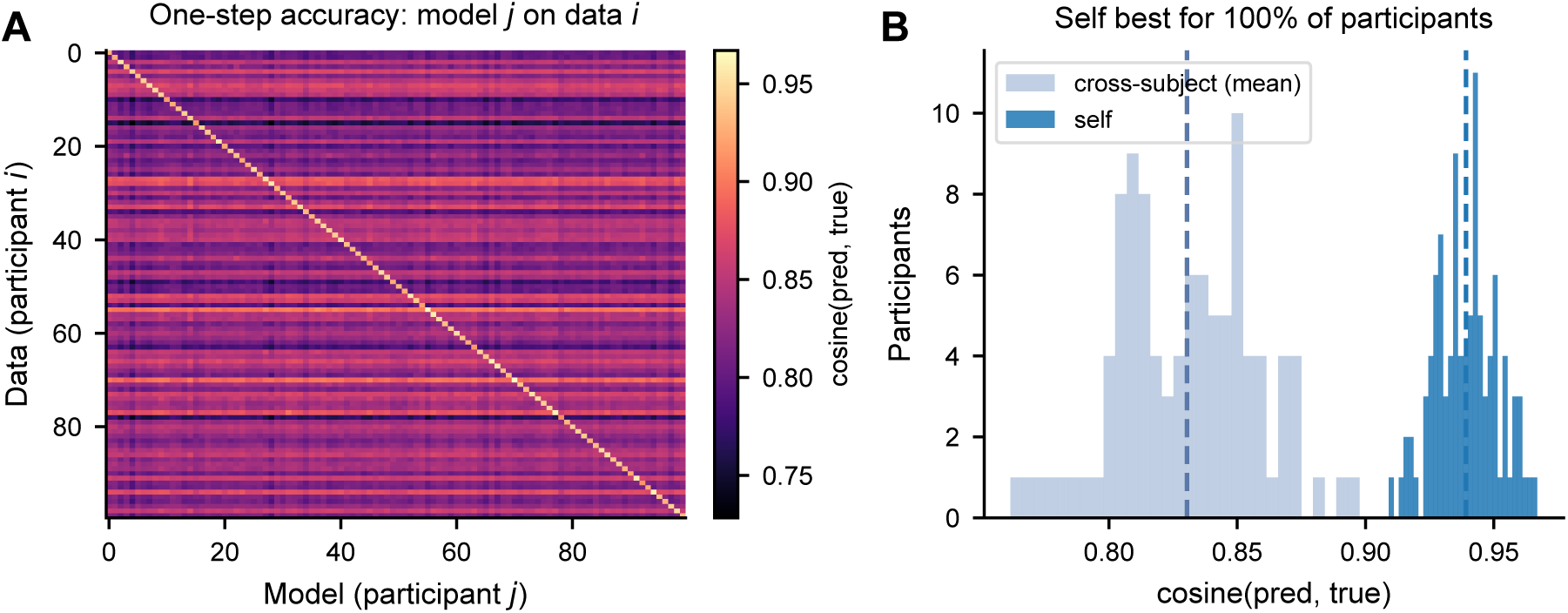
The models are subject-specific: each predicts its own participant better than others. For every participant we held out the last 20% of the recording and measured how well each of the 100 trained models predicts it one step ahead, scored by the cosine similarity between predicted and observed next-state vectors (scale-invariant). **(A)** Accuracy matrix: entry (*i, j*) is the accuracy of model *j* on participant *i*’s held-out data. The bright diagonal shows that each model predicts its own participant best. **(B)** Distribution across participants of the self accuracy (model and data from the same participant) and the mean cross-subject accuracy. Self prediction is higher for every participant (median cosine 0.94 vs 0.83; the participant’s own model is the single best-matching model in 100% of cases; Wilcoxon signed-rank *p <* 10^−17^). The models capture individual dynamics rather than a generic template.

**Figure S5:**
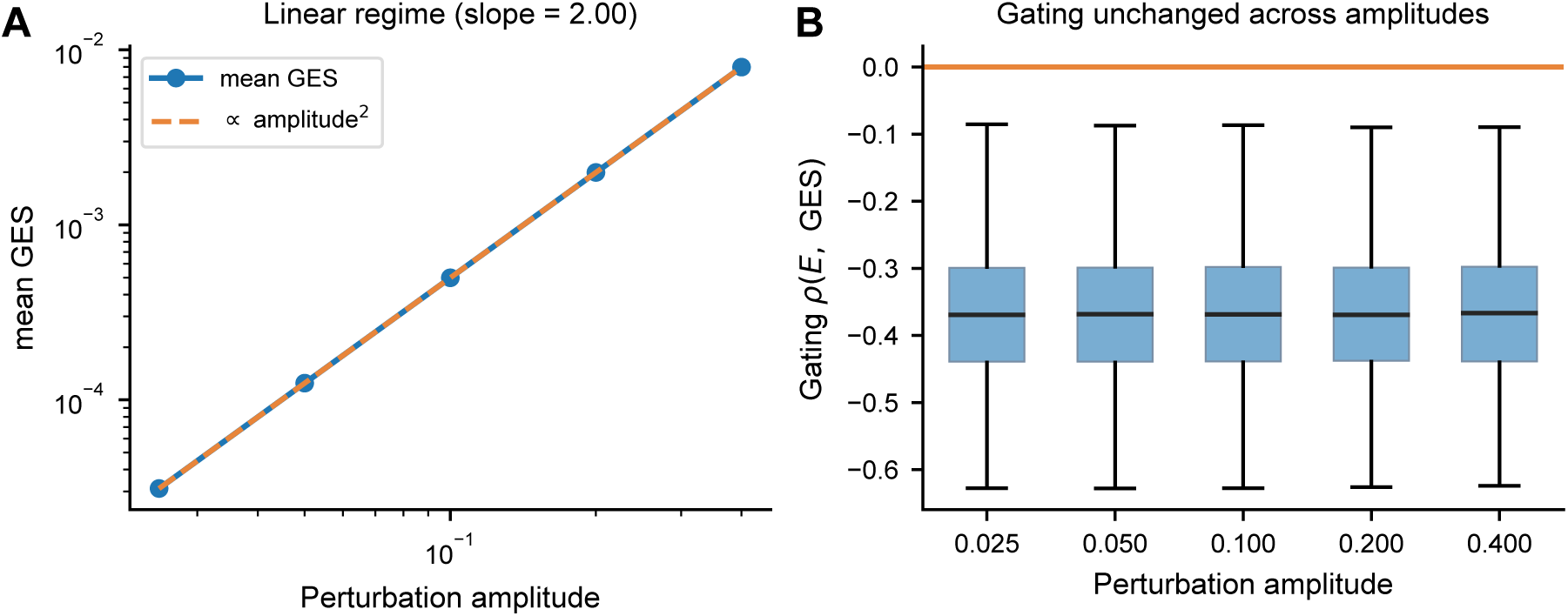
The perturbation analysis is stable across stimulation amplitudes. We repeated the virtual perturbation at amplitudes from 0.025 to 0.4 (in units of the standardized signal; the main analyses use 0.1) and recomputed the global effect size (GES). **(A)** The mean GES scales as the square of the perturbation amplitude (log–log slope = 2.00, dashed guide), confirming that the analysis operates in the linear-response regime where relative effects are amplitude-independent. The spatial map of responsiveness across cortical targets is essentially identical across amplitudes (Spearman *r* = 1.00 against the reference amplitude). **(B)** The state-gating is unchanged: the per-target correlation between baseline energy and GES is the same at every amplitude (median *ρ* ≈ −0.37). Boxplots show the distribution over targets and participants.

**Figure S6:**
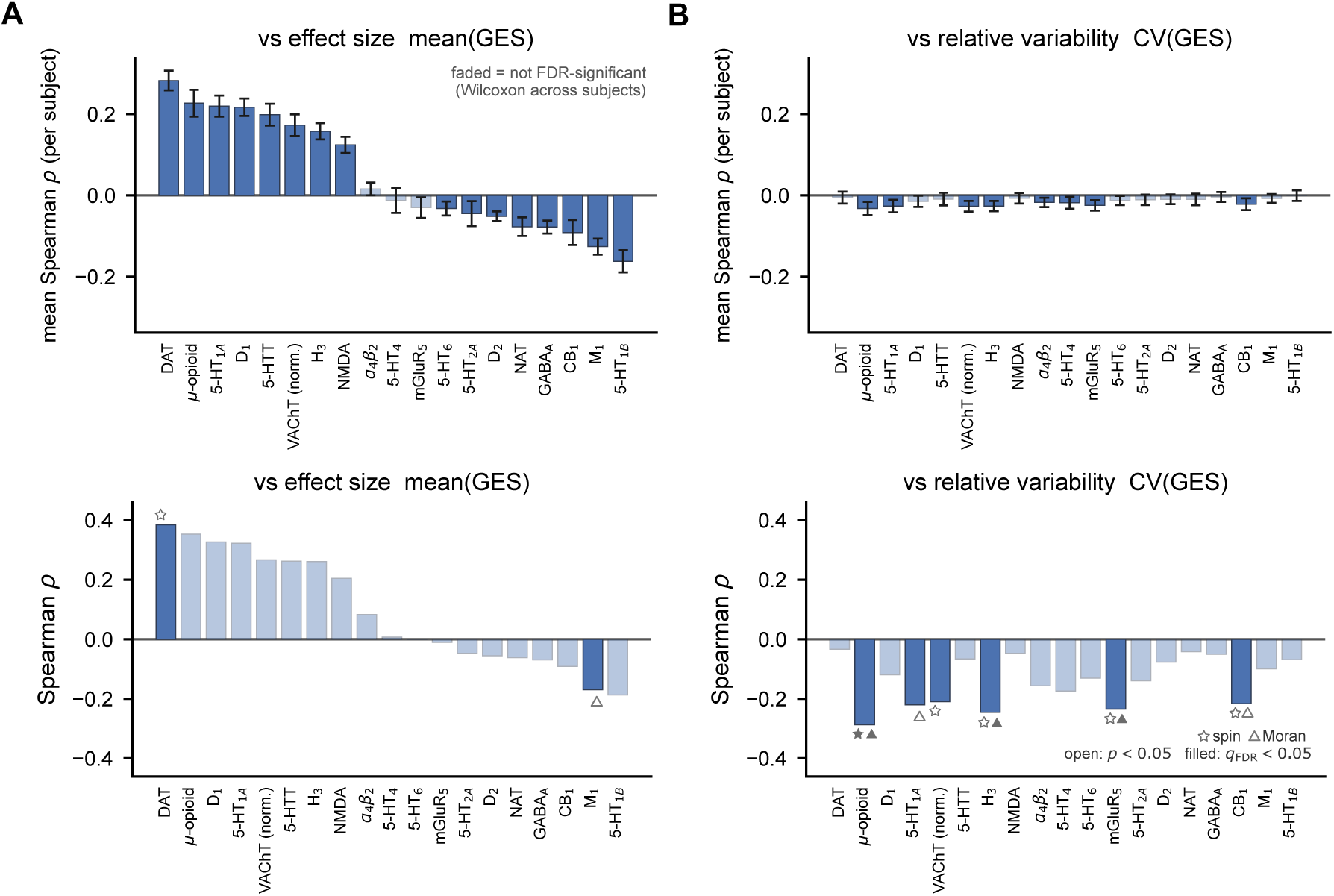
Associations between the regional stimulation response and 19 neurotransmitter receptor and transporter densities. Spearman correlations, across the 400 cortical regions, between each PET-derived density map and two properties of the response: its mean effect size, mean(GES) **(A)**, and its relative trial-to-trial variability, CV(GES) (standard deviation over mean of the effect size across stimulation times) **(B)**. Receptors are ordered by their correlation with mean(GES). *Top row, per-participant analysis:* for each participant, the receptor map is correlated with that participant’s own regional map; bars show the mean correlation across the 100 participants (error bars, 95% confidence interval), and are faded when the across-participant distribution is not significant after FDR correction (Wilcoxon signed-rank versus zero). *Bottom row, group-level analysis:* the receptor map is correlated with the across-participant-mean regional map; significance is assessed against two spatial-autocorrelation-preserving nulls, the spin test (star) and Moran spectral randomization (triangle), with open markers indicating nominal *p <* 0.05 and filled markers survival of the Benjamini–Hochberg FDR correction across the 19 maps (Methods). Response magnitude has no FDR-robust molecular correlate (the dopamine transporter, DAT, is its strongest, nominal under the spin null only), whereas CV(GES) correlates negatively with several neuromodulators (*µ*-opioid, H_3_ and mGluR_5_ survive FDR under the Moran null), i.e. receptor-richer regions respond more reproducibly. These group-level CV associations do not reproduce per participant (top row).

**Figure S7:**
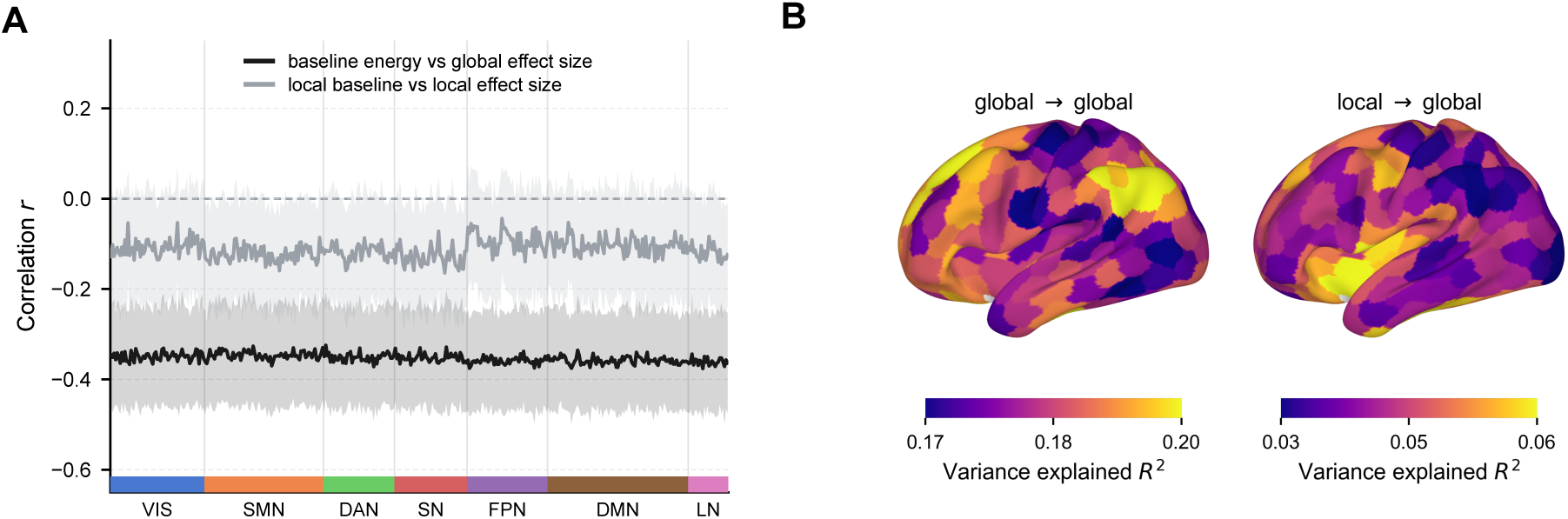
The gating is driven by the whole-brain state, not the target’s local activity. **(A)** For each cortical region (ordered by Yeo-7 network), the across-time correlation between the brain’s ongoing activity before stimulation and the resulting effect size, computed at two scales: the whole-brain baseline energy against the global effect size (black), and the target’s local baseline activity against its local effect size (grey). Lines and shaded bands are the mean ± standard deviation across the 100 participants. The whole-brain correlation is consistently stronger (more negative) than the local one. **(B)** Variance in the global effect size explained (*R*^2^) by the whole-brain baseline energy (global → global, left) and by the target’s local baseline activity (local → global, right), shown on the cortical surface. The whole-brain state explains several-fold more variance than local activity at nearly every region.

**Figure S8:**
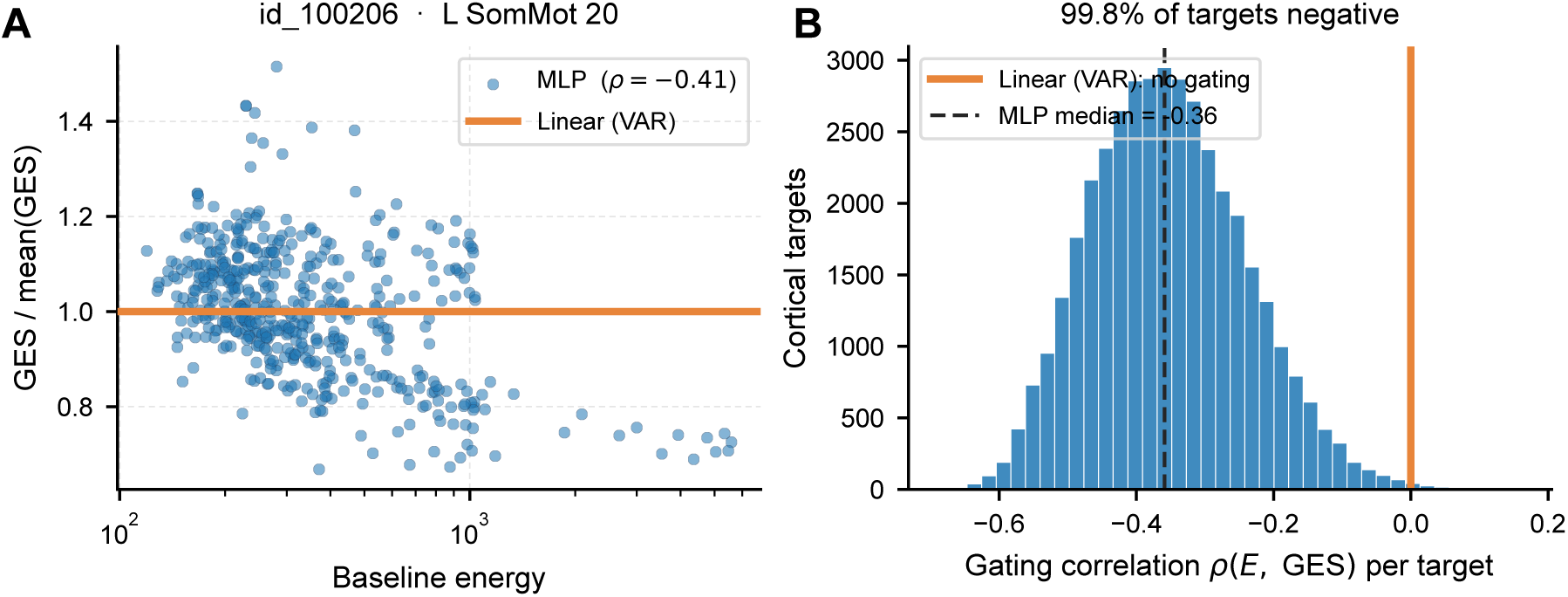
The state-gating is a nonlinear property of the personalized model, absent in a linear model. We fit an optimal linear one-step model (vector-autoregression, ordinary least squares, same *S* = 3 window) to each participant and ran the identical virtual-perturbation and global-effect-size (GES) computation used for the MLP model (Methods). For a linear map the evoked response is independent of the current state, so the GES is constant in time. **(A)** For the example participant and target of Figure 5A (id 100206, L SomMot 20), GES against baseline energy *E*(*t*), normalized to each model’s own temporal mean (the two models have different response magnitudes). The MLP response falls with baseline energy (Spearman *ρ* = −0.41); the linear model is constant (flat line). **(B)** Across cortical targets (100 participants, 40,000 targets), the distribution of the per-target gating correlation *ρ*(*E,* GES) for the MLP (median −0.36, negative in 99.8% of targets). The linear model has zero temporal variance in the GES (coefficient of variation below 10^−13^ at every target and participant), so the correlation is undefined and the gating is absent by construction. The state-dependence therefore reflects nonlinear structure the model learns from the data, not the perturbation or normalization procedure.

**Figure S9:**
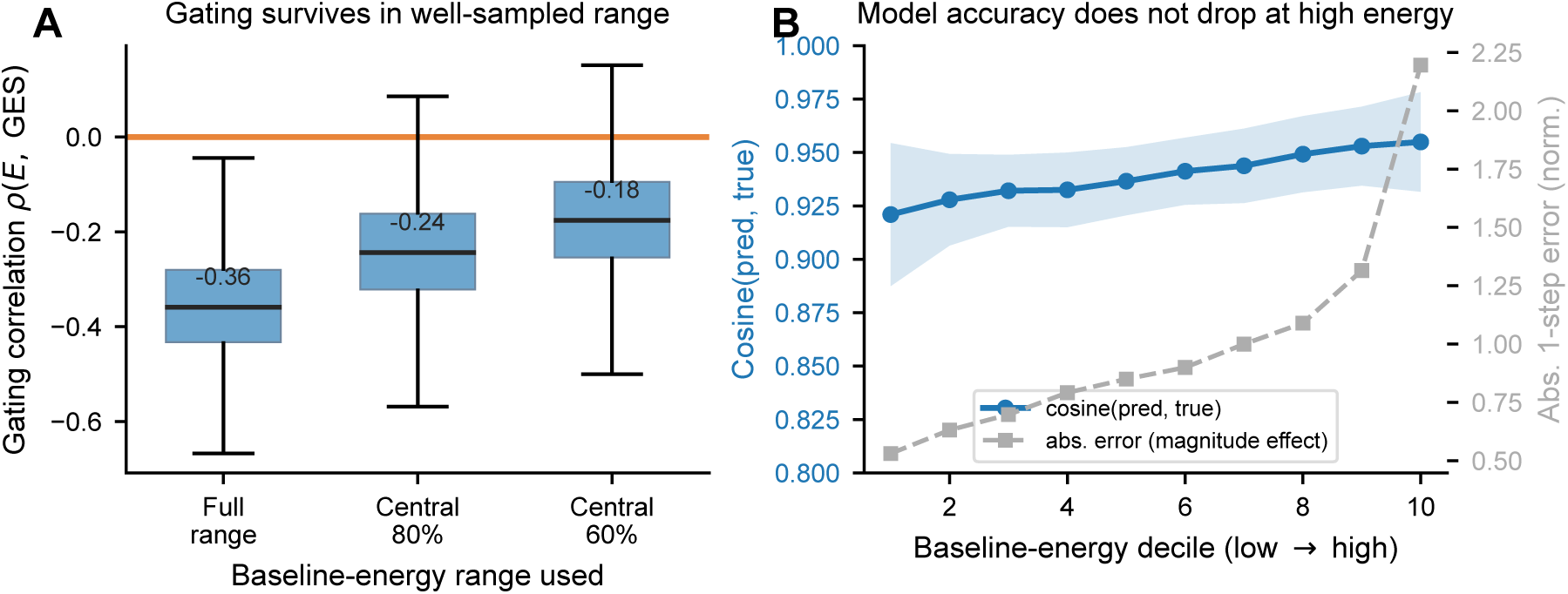
The gating is not an artifact of sparse high-energy sampling. High-energy states are less frequent than low-energy ones, so we checked that the gating is not produced by the model being poorly constrained where data are sparse. **(A)** Per cortical target (100 participants, 40,000 targets), the gating correlation *ρ*(*E,* GES) computed over the full range of baseline energy, over the central 80% ([10,90] percentile), and over the central 60% ([20,80] percentile) of the energy distribution. The gating persists in the densely sampled middle (median *ρ* = −0.18, negative in 92.8% of targets over the central 60%); the attenuation is the expected consequence of restricting the predictor’s range. The orange line marks *ρ* = 0. **(B)** One-step prediction accuracy of the model on held-out data, by baseline-energy decile (low to high). The scale-invariant cosine similarity between predicted and true next-state vectors (blue, left axis) stays high across the whole range and does not fall at high energy (0.92 to 0.96). The normalized absolute error (grey, right axis) rises with energy, but this reflects the larger signal magnitude of high-energy states rather than a loss of accuracy. The model is therefore well-fit across the energy range, so the smaller response at high energy is not a fitting artifact. Lines and bands are mean ± standard deviation across participants.

**Figure S10:**
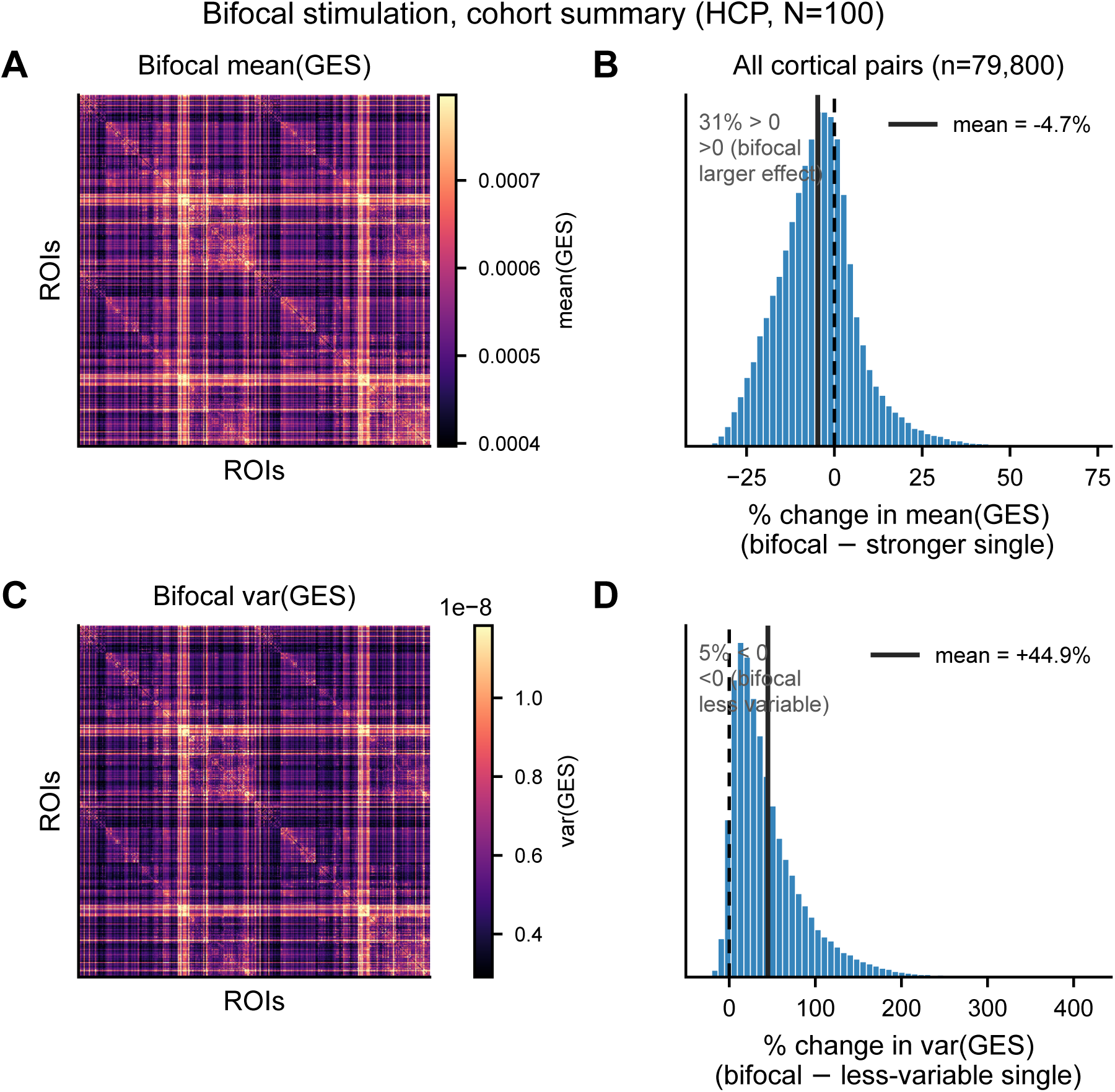
Cohort statistics of bifocal stimulation across all cortical pairs (absolute scale). Group-level summary of the bifocal analysis of Figure 6, over all 79,800 cortical region pairs, averaged across the 100 participants; the network-pair breakdowns are in Figure 6F,G. **(A)** Matrix of the bifocal mean effect size, mean(GES), for every region pair (rows and columns are cortical regions). **(B)** Distribution across all pairs of the percentage change in mean(GES) of bifocal co-stimulation relative to the stronger of the two single sites. Joint stimulation evokes a larger effect than the stronger single site for 31% of pairs (mean change −4.7%); the dashed line marks the mean. **(C)** Matrix of the bifocal absolute variance, var(GES). **(D)** Distribution of the percentage change in var(GES) relative to the less variable of the two single sites. Bifocal stimulation is less variable, in absolute terms, than the less variable single site for only 5% of pairs (mean change +44.9%): because the variance scales with the typically larger joint effect, co-stimulation usually raises the *absolute* variance even though it lowers the *relative* variability (CV) for most pairs (Figure 6C,E,G).

**Table S1:** Symbols and response measures used in the perturbation analysis. For each target region *j*, a virtual perturbation is applied at time *t* and the response is read at *t*+1; the quantities below summarize this response per region, per target, and over stimulation times. mean(GES), Var(GES) and CV(GES) are the response measures used in the main figures; the variance-explained measures are compared in Figure S7.

| Symbol | Definition |
| --- | --- |
| *Indices and states* |  |
| $j$ | stimulus target region (ROI) |
| $x_i(t)$ | baseline activity of region $i$ at time $t$ |
| $x_i(t+1)$ | next unperturbed state of region $i$ |
| $x_i^{(j)}(t+1)$ | next perturbed (evoked) state of region $i$ under stimulation at $j$ |
| *Energy measures* |  |
| $e_i(t) = x_i^2(t)$ | local baseline energy |
| $E(t) = \sum_i x_i^2(t)$ | global baseline energy (total ongoing activity) |
| $e_j^{(j)}(t+1) = (x_j^{(j)}(t+1))^2$ | local evoked energy at the target |
| $E^{(j)}(t+1) = \sum_i (x_i^{(j)}(t+1))^2$ | global evoked energy |
| *Effect size (squared response)* |  |
| $\gamma_j^{(j)} = (x_j^{(j)} - x_j)^2$ | local effect size at the target |
| $\Gamma^{(j)} = \sum_i (x_i^{(j)} - x_i)^2$ | global effect size (GES), total squared response |
| *Effect direction (signed response)* |  |
| $\delta_j^{(j)} = x_j^{(j)} - x_j$ | local effect direction at the target |
| $\Delta^{(j)} = \sum_i (x_i^{(j)} - x_i)$ | global effect direction |
| *Per-target summaries (over stimulation times)* |  |
| $\langle \cdot \rangle_t$ | time-average per target; $\text{mean}(\text{GES}) = \langle \Gamma^{(j)} \rangle_t$ is the responsiveness |
| $\text{Var}_t(\cdot)$ | temporal variance per target; $\text{Var}(\text{GES}) = \text{Var}_t(\Gamma^{(j)})$ is the absolute variability |
| $\text{CV}_t(\cdot)$ | temporal coefficient of variation; $\text{CV}(\text{GES}) = \text{SD}_t(\Gamma^{(j)}) / \langle \Gamma^{(j)} \rangle_t$ is the relative variability |
| $[\text{corr}(a, b)]^2$ | variance in evoked $b$ explained by baseline $a$ (global $E$ or local $e_j$; Fig. S7) |

## References

[1] Massimini M, Ferrarelli F, Huber R, Esser SK, Singh H, Tononi G. Breakdown of cortical effective connectivity during sleep. Science. 2005;309(5744):2228–32.

[2] Deco G, Cabral J, Saenger VM, Boly M, Tagliazucchi E, Laufs H, et al. Perturbation of whole-brain dynamics in silico reveals mechanistic differences between brain states. Neuroimage. 2018;169:46–56.

[3] Momi D, Wang Z, Parmigiani S, Mikulan E, Bastiaens SP, Oveisi MP, et al. Stimulation mapping and whole-brain modeling reveal gradients of excitability and recurrence in cortical networks. Nature Communications. 2025;16(1):3222.

[4] Homan S, Muscat W, Joanlanne A, Marousis N, Cecere G, Hofmann L, et al. Treatment effect variability in brain stimulation across psychiatric disorders: A meta-analysis of variance. Neuroscience & Biobehavioral Reviews. 2021;124:54–62.

[5] Sack AT, Paneva J, Küthe T, Dijkstra E, Zwienenberg L, Arns M, et al. Target engagement and brain state dependence of transcranial magnetic stimulation: implications for clinical practice. Biological Psychiatry. 2024;95(6):536–44.

[6] Briley P, Webster L, Lankappa S, Pszczolkowski S, McAllister-Williams R, Liddle P, et al. Trajectories of improvement with repetitive transcranial magnetic stimulation for treatment-resistant major depression in the BRIGhTMIND trial. npj Mental Health Research. 2024;3(1):32.

[7] Fitzgerald PB, Hoy KE, Anderson RJ, Daskalakis ZJ. A study of the pattern of response to rTMS treatment in depression. Depression and anxiety. 2016;33(8):746–53.

[8] Momi D, Wang Z, Oveisi MP, Kadak K, Bastiaens SP, Lissemore JI, et al. Excitation-Inhibition Balance and Fronto-Limbic Connectivity Drive TMS Treatment Outcomes in Refractory Depression. bioRxiv. 2025:2025–04.

[9] López-Alonso V, Cheeran B, Rio-Rodríguez D, Fernández-del Olmo M. Inter-individual variability in response to non-invasive brain stimulation paradigms. Brain stimulation. 2014;7(3):372–80.

[10] Ziemann U, Siebner HR. Inter-subject and inter-session variability of plasticity induction by non-invasive brain stimulation: boon or bane? Brain Stimulation: Basic, Translational, and Clinical Research in Neuromodulation. 2015;8(3):662–3.

[11] Huang YZ, Lu MK, Antal A, Classen J, Nitsche M, Ziemann U, et al. Plasticity induced by non-invasive transcranial brain stimulation: a position paper. Clinical Neurophysiology. 2017;128(11):2318–29.

[12] Guerra A, López-Alonso V, Cheeran B, Suppa A. Variability in non-invasive brain stimulation studies: Reasons and results. Neuroscience letters. 2020;719:133330.

[13] Jedynak M, Boyer A, Chanteloup-Forêt B, Bhattacharjee M, Saubat C, Tadel F, et al. Variability of single pulse electrical stimulation responses recorded with intracranial electroencephalography in epileptic patients. Brain Topography. 2023;36(1):119–27.

[14] Zrenner C, Ziemann U. Closed-loop brain stimulation. Biological Psychiatry. 2024;95(6):545–52.

[15] Bergmann TO. Brain state-dependent brain stimulation. Frontiers in psychology. 2018;9:2108.

[16] Zrenner C, Desideri D, Belardinelli P, Ziemann U. Real-time EEG-defined excitability states determine efficacy of TMS-induced plasticity in human motor cortex. Brain stimulation. 2018;11(2):374–89.

[17] Desideri D, Zrenner C, Ziemann U, Belardinelli P. Phase of sensorimotor *µ*-oscillation modulates cortical responses to transcranial magnetic stimulation of the human motor cortex. The Journal of physiology. 2019;597(23):5671–86.

[18] Momi D, Ozdemir RA, Tadayon E, Boucher P, Di Domenico A, Fasolo M, et al. Phase-dependent local brain states determine the impact of image-guided transcranial magnetic stimulation on motor network electroencephalographic synchronization. The Journal of physiology. 2022;600(6):1455–71.

[19] Avramiea AE, Hardstone R, Lueckmann JM, Bím J, Mansvelder HD, Linkenkaer-Hansen K. Pre-stimulus phase and amplitude regulation of phase-locked responses are maximized in the critical state. Elife. 2020;9:e53016.

[20] Silvanto J, Muggleton N, Walsh V. State-dependency in brain stimulation studies of perception and cognition. Trends in cognitive sciences. 2008;12(12):447–54.

[21] Bradley C, Nydam AS, Dux PE, Mattingley JB. State-dependent effects of neural stimulation on brain function and cognition. Nature reviews neuroscience. 2022;23(8):459–75.

[22] Marzetti L, Basti A, Chella F, Guidotti R, Pizzella V, Romani GL. Real-time mapping of intrinsic brain connectivity to target brain stimulation. Brain Stimulation: Basic, Translational, and Clinical Research in Neuromodulation. 2021;14(6):1751–2.

[23] Kabir A, Dhami P, Dussault Gomez MA, Blumberger DM, Daskalakis ZJ, Moreno S, et al. Influence of large-scale brain state dynamics on the evoked response to brain stimulation. Journal of Neuroscience. 2024;44(39):e0782242024.

[24] Angiolelli M, Depannemaecker D, Agouram H, Régis J, Carron R, Woodman M, et al. The virtual parkinsonian patient. npj Systems Biology and Applications. 2025;11(1):40.

[25] Rabuffo G, Angiolelli M, Fukai T, Deco G, Sorrentino P, Momi D. Pre-stimulus brain states predict and control variability in stimulation responses. Brain Stimulation. 2026;19(4):103118.

[26] Grosshagauer S, Woletz M, Vasileiadi M, Linhardt D, Nohava L, Schuler AL, et al. Chronometric TMS-fMRI of personalized left dorsolateral prefrontal target reveals state-dependency of subgenual anterior cingulate cortex effects. Molecular Psychiatry. 2024;29(9):2678–88.

[27] Rossi S, Antal A, Bestmann S, Bikson M, Brewer C, Brockmöller J, et al. Safety and recommendations for TMS use in healthy subjects and patient populations, with updates on training, ethical and regulatory issues: Expert Guidelines. Clinical Neurophysiology. 2021;132(1):269–306.

[28] Arieli A, Sterkin A, Grinvald A, Aertsen A. Dynamics of ongoing activity: explanation of the large variability in evoked cortical responses. Science. 1996;273(5283):1868–71.

[29] Bergmann TO, Mölle M, Schmidt MA, Lindner C, Marshall L, Born J, et al. EEG-guided transcranial magnetic stimulation reveals rapid shifts in motor cortical excitability during the human sleep slow oscillation. Journal of Neuroscience. 2012;32(1):243–53.

[30] Massimini M, Rosanova M, Mariotti M. EEG slow ( 1 Hz) waves are associated with nonstationarity of thalamo-cortical sensory processing in the sleeping human. Journal of neurophysiology. 2003;89(3):1205–13.

[31] Polanía R, Nitsche MA, Ruff CC. Studying and modifying brain function with non-invasive brain stimulation. Nature neuroscience. 2018;21(2):174–87.

[32] Deco G, Cruzat J, Cabral J, Tagliazucchi E, Laufs H, Logothetis NK, et al. Awakening: Predicting external stimulation to force transitions between different brain states. Proceedings of the National Academy of Sciences. 2019;116(36):18088–97.

[33] Dagnino PC, Escrichs A, López-González A, Gosseries O, Annen J, Sanz Perl Y, et al. Re-awakening the brain: Forcing transitions in disorders of consciousness by external in silico perturbation. PLOS Computational Biology. 2024;20(5):e1011350.

[34] Sorrentino P, Seguin C, Rucco R, Liparoti M, Lopez ET, Bonavita S, et al. The structural connectome constrains fast brain dynamics. Elife. 2021;10:e67400.

[35] Rabuffo G, Lokossou HA, Li Z, Ziaee-Mehr A, Hashemi M, Quilichini PP, et al. Mapping global brain reconfigurations following local targeted manipulations. Proceedings of the National Academy of Sciences. 2025;122(16):e2405706122.

[36] Acero-Pousa I, Bonetti L, Rosso M, Sanz-Perl Y, Vuust P, Kringelbach ML, et al. Beyond where: When and how brain stimulation drives state transitions. bioRxiv. 2026:2026–03.

[37] Luo Z, Peng K, Liang Z, Cai S, Xu C, Li D, et al. Mapping effective connectivity by virtually perturbing a surrogate brain. Nature Methods. 2025;22:1376–85.

[38] Margulies DS, Ghosh SS, Goulas A, Falkiewicz M, Huntenburg JM, Langs G, et al. Situating the default-mode network along a principal gradient of macroscale cortical organization. Proceedings of the National Academy of Sciences. 2016;113(44):12574–9.

[39] Ferguson KA, Cardin JA. Mechanisms underlying gain modulation in the cortex. Nature Reviews Neuroscience. 2020;21(2):80–92.

[40] Polack PO, Friedman J, Golshani P. Cellular mechanisms of brain state-dependent gain modulation in visual cortex. Nature Neuroscience. 2013;16(9):1331–9.

[41] Hernandez-Pavon JC, Veniero D, Bergmann TO, Belardinelli P, Bortoletto M, Casarotto S, et al. TMS combined with EEG: Recommendations and open issues for data collection and analysis. Brain stimulation. 2023;16(2):567–93.

[42] Burt JB, Demirtaş M, Eckner WJ, Navejar NM, Ji JL, Martin WJ, et al. Hierarchy of transcriptomic specialization across human cortex captured by structural neuroimaging topography. Nature Neuroscience. 2018;21(9):1251–9.

[43] Paquola C, Vos de Wael R, Wagstyl K, Bethlehem RAI, Hong SJ, Seidlitz J, et al. Microstructural and functional gradients are increasingly dissociated in transmodal cortices. PLoS Biology. 2019;17(5):e3000284.

[44] Murray JD, Bernacchia A, Freedman DJ, Romo R, Wallis JD, Cai XJ, et al. A hierarchy of intrinsic timescales across primate cortex. Nature Neuroscience. 2014;17(12):1661–3.

[45] Froudist-Walsh S, Xu T, Niu M, Rapan L, Zhao L, Margulies DS, et al. Gradients of neurotransmitter receptor expression in the macaque cortex. Nature Neuroscience. 2023;26(7):1281–94.

[46] Cocchi L, Sale MV, L Gollo L, Bell PT, Nguyen VT, Zalesky A, et al. A hierarchy of timescales explains distinct effects of local inhibition of primary visual cortex and frontal eye fields. elife. 2016;5:e15252.

[47] Gollo LL, Roberts JA, Cocchi L. Mapping how local perturbations influence systems-level brain dynamics. Neuroimage. 2017;160:97–112.

[48] Sinisalo H, Rissanen I, Kahilakoski OP, Souza VH, Tommila T, Laine M, et al. Modulating brain networks in space and time: Multi-locus transcranial magnetic stimulation. Clinical Neurophysiology. 2024;158:218–24.

[49] Fischer DB, Fried PJ, Ruffini G, Ripolles O, Salvador R, Banus J, et al. Multifocal tDCS targeting the resting state motor network increases cortical excitability beyond traditional tDCS targeting unilateral motor cortex. Neuroimage. 2017;157:34–44.

[50] Blumberger DM, Maller JJ, Thomson L, Mulsant BH, Rajji TK, Maher M, et al. Unilateral and bilateral MRI-targeted repetitive transcranial magnetic stimulation for treatment-resistant depression: a randomized controlled study. Journal of Psychiatry and Neuroscience. 2016;41(4):E58–66.

[51] Kringelbach ML, Perl YS, Deco G. The thermodynamics of mind. Trends in Cognitive Sciences. 2024;28(6):568–81.

[52] Stikvoort W, Pérez-Ordoyo E, Mindlin I, Escrichs A, Sitt JD, Kringelbach ML, et al. Nonequi-librium brain dynamics elicited as the origin of perturbative complexity. PLoS computational biology. 2025;21(6):e1013150.

[53] Berjaga-Buisan T, Monti JM, Cortada M, Colombo MA, Geli SM, Gaglioti G, et al. Thermodynamics of consciousness: A non-invasive perturbational framework. bioRxiv. 2025:2025–12.

[54] Glover GH. Overview of functional magnetic resonance imaging. Neurosurgery Clinics of North America. 2011;22(2):133.

[55] Logothetis NK. What we can do and what we cannot do with fMRI. Nature. 2008;453(7197):869–78.

[56] Siebner HR, Funke K, Aberra AS, Antal A, Bestmann S, Chen R, et al. Transcranial magnetic stimulation of the brain: What is stimulated?–A consensus and critical position paper. Clinical neurophysiology. 2022;140:59–97.

[57] Bergmann TO, Varatheeswaran R, Hanlon CA, Madsen KH, Thielscher A, Siebner HR. Concurrent TMS-fMRI for causal network perturbation and proof of target engagement. Neuroimage. 2021;237:118093.

[58] Rafiei F, Rahnev D. TMS does not increase BOLD activity at the site of stimulation: a review of all concurrent TMS-fMRI studies. eNeuro. 2022;9(4):ENEURO.0163-22.2022.

[59] Van Essen DC, Smith SM, Barch DM, Behrens TEJ, Yacoub E, Ugurbil K. The WU-Minn Human Connectome Project: an overview. NeuroImage. 2013;80:62–79.

[60] Schaefer A, Kong R, Gordon EM, Laumann TO, Zuo XN, Holmes AJ, et al. Local-global parcellation of the human cerebral cortex from intrinsic functional connectivity MRI. Cerebral Cortex. 2018;28(9):3095–114.

[61] Yeo BT, Krienen FM, Sepulcre J, Sabuncu MR, Lashkari D, Hollinshead M, et al. The organization of the human cerebral cortex estimated by intrinsic functional connectivity. Journal of neurophysiology. 2011.

[62] Tian Y, Margulies DS, Breakspear M, Zalesky A. Topographic organization of the human sub-cortex unveiled with functional connectivity gradients. Nature Neuroscience. 2020;23(11):1421–32.

[63] Salimi-Khorshidi G, Douaud G, Beckmann CF, Glasser MF, Griffanti L, Smith SM. Automatic denoising of functional MRI data: combining independent component analysis and hierarchical fusion of classifiers. NeuroImage. 2014;90:449–68.

[64] Faskowitz J, Esfahlani FZ, Jo Y, Sporns O, Betzel RF. Edge-centric functional network representations of human cerebral cortex reveal overlapping system-level architecture. Nature Neuroscience. 2020;23(12):1644–54.

[65] Esfahlani FZ, Jo Y, Faskowitz J, Byrge L, Kennedy DP, Sporns O, et al. High-amplitude cofluctuations in cortical activity drive functional connectivity. Proceedings of the National Academy of Sciences. 2020;117(45):28393–401.

[66] Rabuffo G, Fousek J, Bernard C, Jirsa V. Neuronal cascades shape whole-brain functional dynamics at rest. ENeuro. 2021;8(5).

[67] Hansen JY, Shafiei G, Markello RD, Smart K, Cox SML, Nørgaard M, et al. Mapping neurotransmitter systems to the structural and functional organization of the human neocortex. Nature Neuroscience. 2022;25(11):1569–81.

[68] Markello RD, Hansen JY, Liu ZQ, Bazinet V, Shafiei G, Suárez LE, et al. neuromaps: structural and functional interpretation of brain maps. Nature Methods. 2022;19(11):1472–9.

[69] Barch DM, Burgess GC, Harms MP, Petersen SE, Schlaggar BL, Corbetta M, et al. Function in the human connectome: task-fMRI and individual differences in behavior. NeuroImage. 2013;80:169–89.

[70] Alexander-Bloch AF, Shou H, Liu S, Satterthwaite TD, Glahn DC, Shinohara RT, et al. On testing for spatial correspondence between maps of human brain structure and function. NeuroImage. 2018;178:540–51.

[71] Wagner HH, Dray S. Generating spatially constrained null models for irregularly spaced data using M oran spectral randomization methods. Methods in Ecology and Evolution. 2015;6(10):1169–78.

[72] Benjamini Y, Hochberg Y. Controlling the false discovery rate: a practical and powerful approach to multiple testing. Journal of the Royal statistical society: series B (Methodological). 1995;57(1):289–300.

